# Dysregulation of the fluid homeostasis system by aging

**DOI:** 10.1101/2024.09.26.615271

**Authors:** Heeun Jang, Alexis Behne Sharma, Usan Dan, Jasmine H. Wong, Zachary A. Knight, Jennifer L. Garrison

**Author notes:** Department of Biomedical Sciences, College of Veterinary Medicine, Iowa State University, Ames, IA 50011, USA.

## Abstract

Chronic dehydration is a leading cause of morbidity for the elderly, but how aging alters the fluid homeostasis system is not well understood. Here, we used a combination of physiologic, behavioral and circuit analyses to characterize how fluid balance is affected by aging in mice. We found that old mice have a primary defect in sensing and producing the anti-diuretic hormone vasopressin, which results in chronic dehydration. Recordings and manipulations of the thirst circuitry revealed that old mice retain the ability to sense systemic cues of dehydration but are impaired in detecting presystemic, likely oropharyngeal, cues generated during eating and drinking, resulting in disorganized drinking behavior on short timescales. Surprisingly, old mice had increased drinking and motivation after 24-hour water deprivation, indicating that aging does not result in a general impairment in the thirst circuit. These findings reveal how a homeostatic system undergoes coordinated changes during aging.

## BACKGROUND

Tight control of fluid balance is essential for life. This is achieved by a physiologic system that monitors the osmolality and volume of the blood and, in response to dehydration, triggers two counterregulatory responses: water consumption, which is motivated by the sensation of thirst, and water reabsorption by the kidney, which is triggered by the hormone vasopressin (AVP). These two responses are controlled by dedicated neural circuits in the forebrain that directly sense changes in fluid balance (reviewed in ^1–4^). Two key nodes in this forebrain circuitry are the subfornical organ (SFO), which contains glutamatergic neurons (SFO^Glut^ neurons) that are activated by increases in blood osmolality and decreases in blood volume to produce thirst (**Fig. 1a**), and AVP-expressing neurons in the supraoptic and paraventricular hypothalamic (PVH) nuclei, which produce and release AVP into the blood in response to upstream dehydration signals, including those arising from the SFO (**Fig. 1b**). These two nodes are interconnected with other forebrain nuclei that sense and process fluid balance information, including the organum vasculosum of the lamina terminalis (OVLT) and the median preoptic nucleus (MnPO; **Fig. 1a, b**).

**Figure 1.**
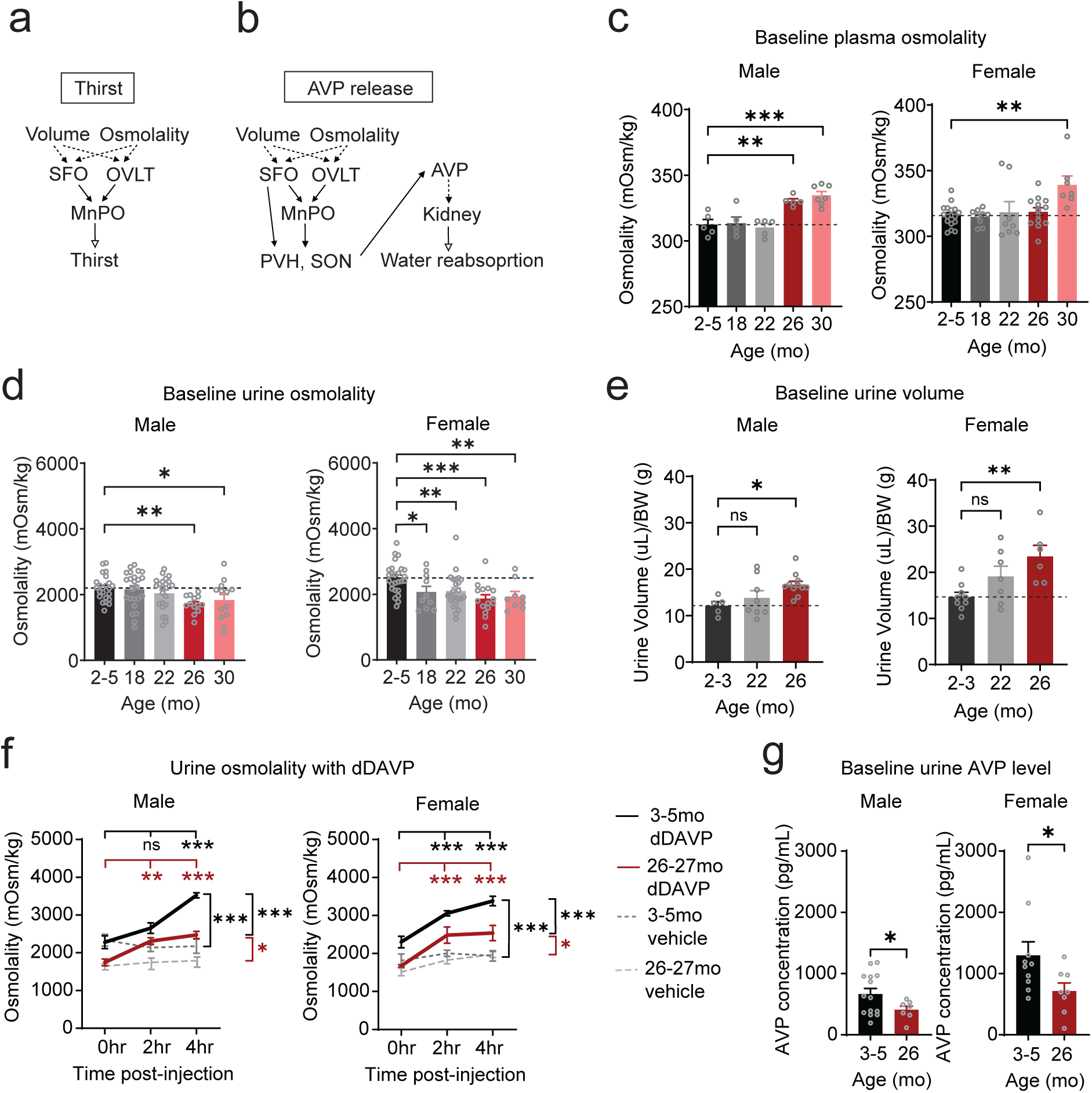
Aged mice are chronically dehydrated and have renal dysfunction. **a.** Schematic of the neural pathway for thirst. Volume and osmolality changes are sensed by the subfornical organ (SFO) and the organum vasculosum of the lamina terminalis (OVLT) and these signals are relayed to their downstream nuclei, including the median preoptic nucleus (MnPO). **b.** Schematic of the neural pathway for AVP release. Volume and osmolality changes are sensed by the SFO and relayed to the MnPO. Both SFO and MnPO project to the paraventricular (PVH) and supraoptic (SON) hypothalamus where AVP is released into circulation to promote water reabsorption in the kidney. In **a** and **b,** solid arrows denote neural innervations, dotted arrows denote sensing of circulatory signals and open arrows indicate functional outcomes **c.** Plasma osmolality in 2-5, 18-19, 22-23, 26-27 and 30-31 month-old male and female mice in their baseline state with free access to chow and water. Male, ***P<0.001 and female, **P<0.01 by one-way ANOVA.**P<0.01 and ***P<0.001 by Fisher’s LSD test. **d.** Urine osmolality in 2-5, 18-19, 22-23, 26-27 and 30-31 month-old male and female mice in their baseline state with *ad libitum* access to chow and water. Male, *P<0.05 and female, ***P<0.001 by one-way ANOVA. *P<0.05, **P<0.01 and ***P<0.001 by Fisher’s LSD test. **e.** Urine volume for 4 hours in 2-3, 22-23 and 26-27 month-old male and female mice adjusted by body weight in 2-3, 22-23 and 26-27 months old male and female mice. Male, *P<0.05 and female, **P<0.01 by one-way ANOVA. ns, non-significant; *P<0.05, **P<0.01 by Dunnet’s test. **f.** Urine osmolality in 3-5 and 26-27 month-old male and female mice before and after dDAVP (1 mg/kg BW) or vehicle injection. For males, ***P<0.001 for main effects of dDAVP [F(2, 80)=14.78] and age [F(3, 80)=34.57], and ***P<0.001 for interactions of their main effects [F(6, 80)=7.240] by 2-way ANOVA. For females, ***P<0.001 for main effects of dDAVP [F(2, 64)=23.50] and age [F(3, 64)=49.55], and *P<0.05 for interactions of their main effects [F(6, 64)=2.775] by 2-way ANOVA. ns, non-significant; *P<0.05 and ***P<0.001 compared to the baseline value or by Dunnet’s test. Values at 4 hours post-dDAVP and post-vehicle within the age group were compared (*P<0.05 and ***P<0.001 by Sidak’s test). Values at 4 hours post-dDAVP were compared between age groups (male, ***P<0.001 and female, ***P<0.001 by Sidak’s test). **g.** Urine AVP levels in 3-5 and 26-27 months-old male and female mice in the baseline state. *P<0.05 by student’s t-test.

Dysregulation of fluid homeostasis is a common feature of aging. For example, older adults report a reduced perception of thirst and consume less water after many thirst-evoking (dipsogenic) stimuli^5–12^. In addition, the ability of the kidney to concentrate urine declines with age, leading to greater loss of fluid in older adults (reviewed in ^13^). As a result, aging is associated with increased prevalence of chronic dehydration^14^, which is a significant risk factor for morbidity and mortality^15,16^.

The specific alterations in the fluid homeostasis system that are caused by aging are not well understood. One challenge is that fluid balance involves multiple interacting systems, including a neuroendocrine system that controls water resorption (the AVP-kidney axis); a sensory system that monitors fluid balance and ingestion, which includes SFO^Glut^ neurons and their sensory afferents arising from the mouth, throat and viscera; and a motivational system which drives water seeking and consumption, which includes SFO^Glut^ neurons as well as their downstream targets such as the dopamine system. It has only recently become possible to monitor and manipulate these fluid homeostasis neurons in behaving animals^17–23^, and no study has examined how any of these circuit nodes are altered by the aging process at the level of specific neural cell types, dynamics, and interconnections.

Here we have performed a comprehensive analysis of how the fluid homeostasis system is altered by aging in mice. We investigated animals of both sexes, across a range of ages from young to very old, and subjected them to batter of analyses at different levels, including: (1) physiologic measurements of fluid balance, kidney function, and AVP release and sensitivity; (2) behavioral analyses of drinking and motivation in response to diverse thirst stimuli (food, dehydration, hyperosmolality, and hypovolemia); (3) neural recordings from circuit nodes that control drinking (SFO^Glut^), AVP release (PVH^AVP^ neurons), and motivation (dopamine release in nucleus accumbens), and (4) optogenetic manipulations to test the sufficiency of circuit nodes. These experiments revealed that a subset of these functions is impaired during aging, whereas others are unexpectedly enhanced. These findings provide for the first time a comprehensive view of how a homeostatic system is altered by the aging process and lay the groundwork for targeted interventions to combat dehydration in the elderly.

## RESULTS

### Aged mice are chronically dehydrated and have renal dysfunction

We set out to characterize how aging affects fluid homeostasis in the mouse. We first measured plasma osmolality in mice of different ages (spanning 2 to 30 months) that had *ad libitum* access to food and water. We found that plasma osmolality was elevated in male mice beginning at 26 months of age (312 ± 4 vs. 330 ± 2 mOsm/kg, 2-5 mo. vs. 26 mo., P = 0.005, **Fig 1c**) and in female mice at 30 months of age (318 ± 3 vs. 339 ± 7 mOsm/kg, 2-5 mo. vs. 26 mo., P < 0.001, **Fig 1c**). Of note, 26-30 months in the mouse corresponds to 73 - 81 years of age in humans^24^, indicating that in mice, as in humans, aging is associated with chronic dehydration.

Fluid homeostasis is governed by two primary mechanisms: water consumption (drinking) and water loss from the kidney. We first characterized renal function in aging mice. We found that aging was associated with reduced urine osmolality, indicating a potential defect in water resorption, in both male (2201 ± 80 vs. 1716 ± 74 mOsm/kg, 2-5 mo. vs. 26 mo., P = 0.0042, **Fig 1d**) and female mice (2500 ± 90 vs. 1864 ± 123 mOsm/kg, 2-5 mo. vs. 26 mo., P = 0.0001, **Fig 1d**). Consistently, urine volume was increased with age in male (12.3 ± 0.9 vs. 16.7 ± 0.7 µL/g BW, 2-3 mo. vs. 26 mo., P = 0.0134, **Fig 1e**) and female mice (14.7 ± 0.7 vs. 23.5 ± 2.4 µL/g, 2-5 mo. vs. 26 mo., P = 0.0028, **Fig 1e**). This suggests that the dehydration in aging mice may be caused, in part, by decreased renal function.

Increased urine volume could be caused by a deficiency in the hormone vasopressin (AVP), which controls renal water absorption. Because it is technically challenging to measure AVP levels directly in old mice (due, in part, to the large blood volume required) we used two surrogate assays. First, we measured AVP concentration in the urine (which directly correlates with plasma AVP levels^25,26^) and found that AVP levels were significantly lower in aged mice (male: 665 ± 91 vs. 409 ± 60 pg/mL, 3-5 mo. vs. 26 mo., P = 0.0301; female: 1296 ± 222 vs. 715 ± 131 pg/mL 3-5 mo. vs. 26 mo., P = 0.0404, **Fig 1g** and **Extended Data Fig 3c**). Second, we assessed plasma copeptin, which is produced as part of the same precursor peptide as AVP, preprovasopressin^27,28^, and is a stable and widely accepted surrogate marker for AVP levels in humans^29–31^ and in mice^32^. We found that there was a trend toward reduced copeptin levels in old mice, although this was not significant (**Extended Data Fig 1a)**. Of note, because old mice are chronically dehydrated (**Fig. 1c**) they should have *elevated* AVP levels relative to young controls^30,33^. The fact that this was not observed suggests that a relative AVP deficiency contributes to their decreased function.

To determine whether there is also AVP resistance in old animals, we challenged mice with a high dose of desmopressin (dDAVP), a selective agonist of the AVP V2 receptor that is expressed in the kidney^34,35^, thereby maximally activating renal AVP signaling. We found that dDAVP significantly increased urine osmolality (reflecting enhanced urine concentrating ability) in both young and aged mice (**Fig 1f** and **Extended Data Fig 1b**). Although aged mice exhibited a relative increase in urine osmolality comparable to that of young mice (**Extended Data Fig 1b**), their absolute urine osmolality remained lower than that of young controls four hours after dDAVP treatment (male: 3523 ± 68 vs. 2389 ± 117 mOsm/kg, young vs. aged, P < 0.001; female: 3382 ± 126 vs. 2539 ± 201 mOsm/kg, young vs. aged, P < 0.001, **Fig 1f**), indicating a relative resistance to AVP. Taken together, these data show that aged mice are chronically dehydrated and that this involves a combination of AVP deficiency and resistance.

### Aging alters the response of AVP neurons to eating and drinking

AVP is produced by neurosecretory cells located in the paraventricular hypothalamus (PVH) and supraoptic nucleus (SON). Given that circulating AVP levels are reduced with aging in mice (**Fig 1g**), we investigated whether the activity of PVH^AVP^ neurons during eating and drinking is also altered in old animals.

We prepared mice for fiber photometry recordings of PVH^AVP^ neurons by injecting a Cre-dependent AAV expressing the calcium indicator GCaMP6s into the PVH of AVP^Cre/+^ male mice and, in the same surgery, implanted an optical fiber above the PVH (**Fig 2a, b**). Because consumption of dry food is a primary stimulus for AVP secretion in rodents^22,36^, we measured AVP neuron responses to food consumption. Mice were fasted overnight and then provided dry chow for 30 min (without water) followed by water access for 30 min without chow (**Fig. 2c**). We found that PVH^AVP^ neurons were gradually activated during chow consumption in both young and old mice^22,36^ (**Fig. 2d**). However, this AVP activation was significantly slower in old male mice compared to young controls (time to half max, 13.7 ± 1.0 vs. 18.3 ± 0.9 min, young vs. aged, P = 0.006, **Fig 2e**) and the magnitude of activation over the entire trial was reduced (mean z score: 1.06 ± 0.11 vs. 0.76 ± 0.07 z, young vs. aged, P = 0.04, **Fig 2g**). Moreover, examination of the microstructure of feeding revealed that young mice exhibited spikes in AVP activity after feeding bouts terminated, while these spikes were reduced in old animals (**Fig. 2i, j**). These differences in calcium dynamics are unlikely to be due to differences in behavior, because there was no difference between young and old animals in the latency to initiate feeding (36.3 ± 6.4 vs. 37.5 ± 3.1 sec, young vs. aged, P = 0.87, **Fig 2f**) or in the amount of chow consumed during the trial (0.37 ± 0.02 vs. 0.37 ± 0.06 g, young vs. aged, P = 0.97, **Fig 2h**). Thus, aged male mice show decreases in AVP neuronal responses to food.

**Figure 2.**
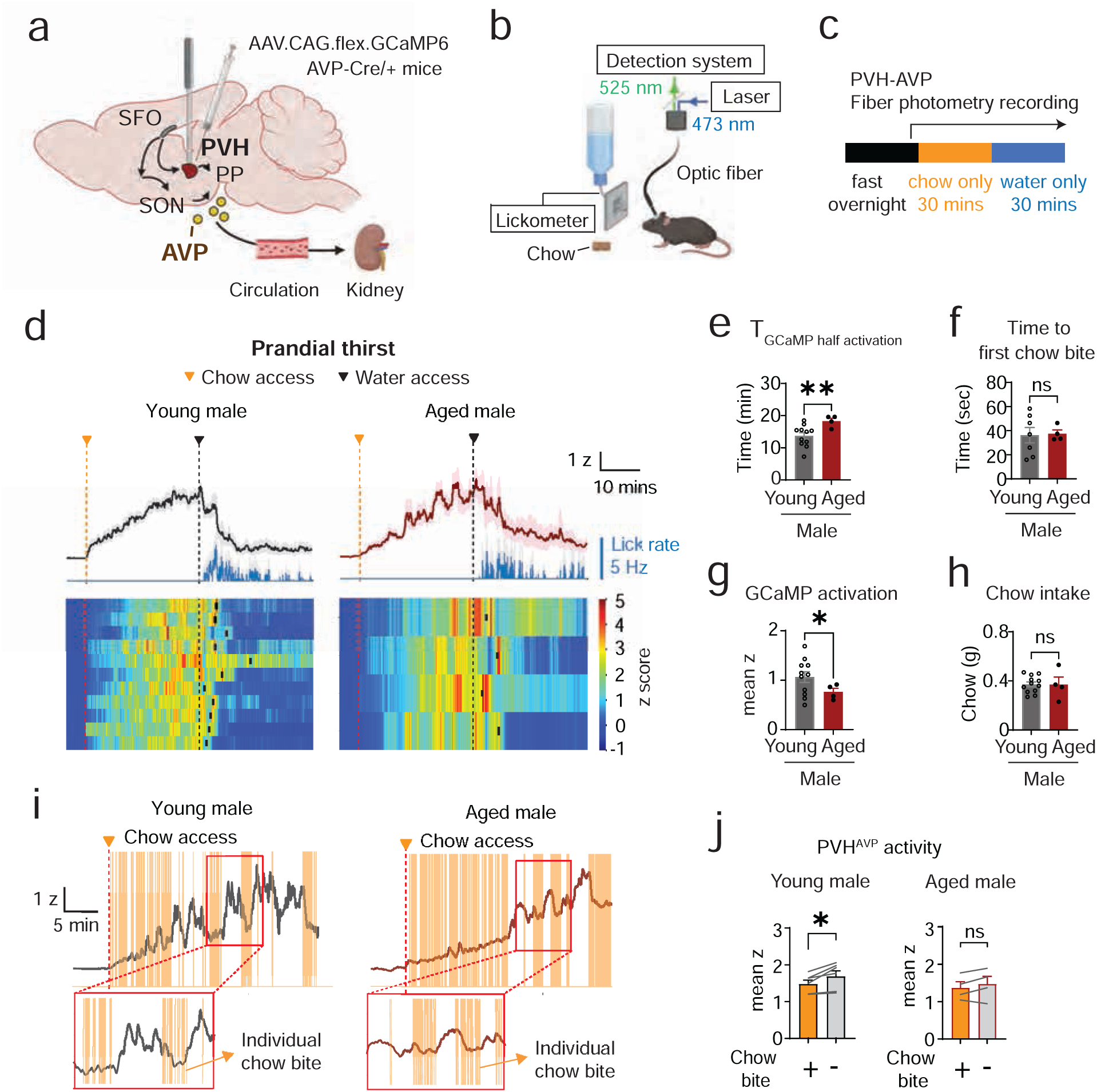
Age delays PVH^AVP^ neuronal responses to feeding. **a.** Schematic for the stereotaxic injection of AAVs cre-dependently expressing GCaMP into the PVH and implantation of an optic fiber for fiber photometry. AVP, arginine vasopressin. PVH, paraventricular nucleus of the hypothalamus, PP, posterior pituitary. SON, supraoptic nucleus. SFO, subfornical organ. Arrows denote synaptic projections. **b.** Schematic for PVH^AVP^ fiber photometry recording during dry chow intake followed by water drinking. **c.** Experimental paradigm for PVH^AVP^ fiber photometry during 30 minutes chow intake after overnight fasting followed by 30-minute water intake. **d.** (Top) Mean lick rate (blue) and mean z-score trace of GCaMP fluorescence in young (3-4 mo, black) and aged (24-26 mo, magenta) male mice, aligned by chow access (red dotted line). (Bottom) Heatmaps of individual z-score traces. Black dotted lines indicate water access. Black rectangles mark the onset of water lick bout in each recording. **e.** Time for GCaMP signals to rise from baseline to the half peak fluorescence. **P<0.01 by Welch’s t-test. **f.** Latency from chow access to the first chow bite. ns., non-significant by Welch’s t-test. **g.** Mean z-score of GCaMP fluorescence from chow access (t=0, red dotted line in **d**) to water access (t=30 mins, black dotted line in **d**). *P<0.05 by Welch’s t-test. **h.** Chow intake during 30 minutes of chow access. ns., non-significant by Welch’s t-test. **i.** Representative PVH^AVP^ z-score traces during chow consumption without water following overnight fasting. Red dotted lines indicate chow access, and yellow vertical bars denote individual chow bite events. Example recordings are shown from 4- (left) and 26-month-old (right) male mice. Red boxes highlight magnified views of GCaMP traces and bite events. **j.** Mean z-score during chow bite (on chow) and between chow bites (off chow) in young and aged male mice. **P<0.001 and *P<0.05 for main effects of chow bite [F(1, 18)=28.81] and age [F(1, 18)=5.316], respectively, and **P<0.01 for their interactions [F(1, 18)=8.899] by 2-way ANOVA. ns., non-significant and ***P<0.001 by Sidak’s test.

AVP neurons are rapidly inhibited by ingestion of water^22,36^. When given water after food, young and old mice both engaged in drinking (**Fig 2d** and **Extended Data Fig 2b-d**) and PVH^AVP^ neurons were inactivated to a similar extent (-1.8 ± 0.3 vs. -1.6 ± 0.4 z, young vs. aged, P = 0.62, **Extended Data Fig 2e**). This inactivation was slower in aged compared to young controls (1.3 ± 0.2 vs. 2.5 ± 0.5 min, young vs. aged, P = 0.01, **Extended Data Fig 2f**), but there was also a trend toward slower water intake in aged animals (mean lick rate: 2.9 ± 0.5 vs. 1.8 ± 0.9 Hz, young vs. aged, P = 0.11, **Extended Data Fig 2g**), which potentially confounds interpretation of neural activity differences. To better understand this drinking phenotype, we investigated next how water intake is altered with age.

### Drinking after eating bouts is delayed in aged mice

Rodents, like humans, consume most of their water during meals^37–41^. We therefore investigated the dynamics of eating and drinking in aged mice with *ad libitum* access to dry food and water (**Fig 3a**). Because prior studies of eating and drinking in old rodents had limited temporal resolution and were only performed in males^42–44^, we used an automated food dispenser and lickometer system to record intake behavior over days at subsecond resolution in both male and female mice at young (3-5 m.o.) and old (26-27 m.o.) ages.

**Figure 3.**
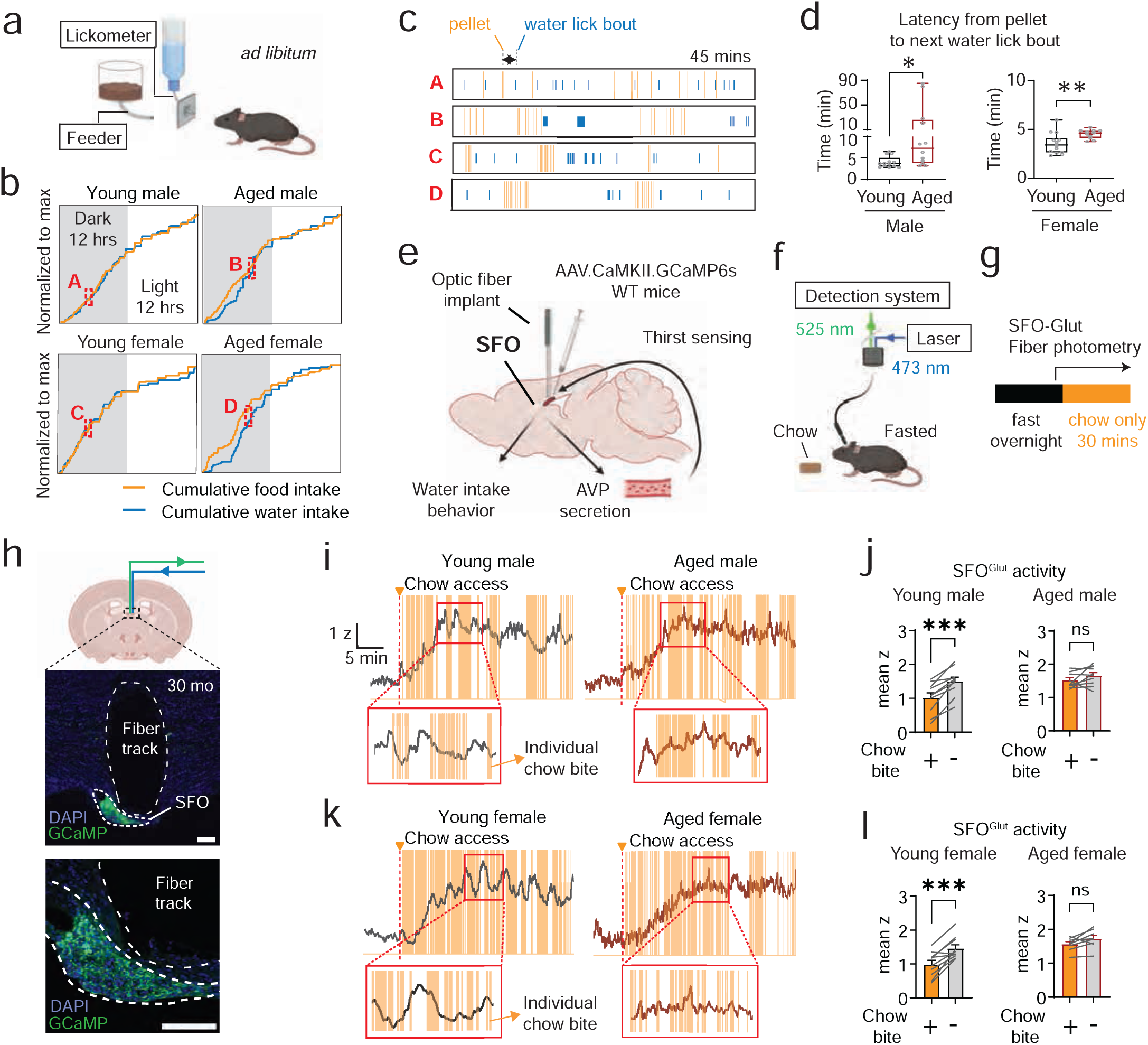
Age delays feeding-induced water drinking. **a.** Schematic for *ad libitum* food and water intake behavior. **b.** Individual examples of cumulative food and water intake over 24 hours, normalized to total food and water intake, respectively, in 3 month-old and 27 months-old male and female mice. The first 12 hours correspond to the dark cycle (gray shaded area), followed by 12 hours of light cycle. **c.** 45 minutes of zoomed-in raster plots from gray and red dotted boxes in **b**; brown bars represent individual pellet-retrieval events and blue bars represent individual water-lick events. **d.** Latency from each pellet retrieval to the onset of the subsequent water-lick bout. Each dot represents the median latency across 24 hours for an individual mouse within each sex and age group. *P<0.05 and **P<0.01 by unpaired t-test. **e.** Schematic for stereotaxic AAV delivery to express GCaMP in the SFO and implantation of an optic fiber for fiber photometry. The SFO senses circulating thirst signals and regulates water intake and AVP secretion. **f.** Schematic for SFO^Glut^ fiber photometry recording during dry chow intake. **g.** Experimental paradigm for SFO^Glut^ fiber photometry recording during 30 minutes of chow intake following overnight fasting. **h.** (Top) Coronal section of the mouse brain indicating the SFO (dotted box). (Middle) GCaMP expression (green) in the SFO and the fiber track (dotted outline) of a 30-month-old male mouse. (Bottom) Zoomed-in image of GCaMP expression in the SFO and fiber track. Scale bars, 100 µm. **i.** Individual z-score traces of GCaMP fluorescence from SFO fiber photometry recording during feeding chow without water after overnight fasting. Dotted red lines represent chow access. Yellow solid vertical bars represent individual chow bite events. Example traces are from (left) 3- and (right) 26-month-old male mice. Red boxes show zoomed in GCaMP trace and individual chow bite events. **j.** Mean z score during chow bite and between chow bites in young and aged male mice. ns, non-significant and ***P<0.001 by paired t-test. **k.** Individual z-score traces of GCaMP fluorescence from SFO^Glut^ fiber photometry recording during feeding chow without water after overnight fasting. Dotted red lines represent chow access. Yellow solid vertical bars represent individual chow bite events. Example traces are from (left) 4- and (right) 27-month-old female mice. Red boxes show zoomed in GCaMP trace and individual chow bite events. **l.** Mean z-score during chow bite and between chow bites in young and aged female mice. ns, non-significant and ***P<0.001 by paired t-test.

We observed no significant difference in the total amount of food and water consumed per day in young and old male mice (**Extended Data Fig 3a**). Because water intake is highly sensitive to the amount of dry food consumed, we also normalized water intake to the amount of food intake, which also showed no significant age-related changes on a timescale of 24 hours (**Extended Data Fig 3b**). This indicates there are no gross changes in total water consumption per day in aged animals. However, it is important to note that because aged animals are chronically dehydrated (**Fig 1c**), which is due, in part, to a failure to appropriately concentrate their urine and retain water (**Fig 1d, e**), this unchanged level of intake represents a failure to increase drinking behavior to match physiologic need.

We wondered whether young and old animals might show differences in their temporal patterns of eating and drinking. To probe this, we analyzed the relationship between consumption of food pellets and the initiation of subsequent drinking (**Fig 3a, b**). We found that young mice exhibited a tight temporal coupling between food and water intake, whereas in aged mice the latency between pellet retrieval and the subsequent lick bout was significantly increased (male: 4.0 ± 0.5 vs. 21.3 ± 14.0 min, young vs. aged, P = 0.008; female: 3.5 ± 0.4 vs. 4.5 ± 0.2 min, young vs. aged, P = 0.0062, **Fig 3c, d** and **Extended Data Fig 3c**). Importantly, this delay was not due to slower pellet consumption in aged animals. In fact, the interval between successive pellet retrievals within meals of comparable size was similar in aged male mice and even shortened in aged female mice compared to young controls (**Extended Data Fig 3f, h**). This reflects that aged mice often retrieved more pellets and spilled a substantial fraction, thereby taking longer to finish a similar size of meal (**Extended Data Fig 3g**). However, the initiation of drinking bout after the last pellet of a meal was delayed in aged mice (male: 4.5 ± 0.8 vs. 12.3 ± 4.5 min, young vs. aged, P = 0.0737; female: 3.1 ± 0.3 vs. 4.4 ± 0.2 min, young vs. aged, P = 0.0017, **Extended Data Fig 3i**), further supporting a specific impairment in feeding-induced thirst. Of note, when aged mice did initiate drinking, they more frequently exhibited larger and longer lick bouts than young animals, possibly as a compensatory response to delayed bout initiation (**Extended Data Fig 3k-n**). Thus, aged mice have a specific defect in the short-timescale temporal coupling between eating and drinking.

### Food intake and SFO activity are uncoupled on short timescales in aged mice

The age-related delay in water consumption following food ingestion could be due to (1) a delayed activation of primary thirst circuits or (2) delayed responses in downstream cells. To distinguish between these possibilities, we monitored the activity of SFO^Glut^ neurons, which are primary sensory neurons that are activated by food consumption and promote thirst^1,4,17,21,45^. We prepared mice for fiber photometry recordings of SFO^Glut^ neurons (**Fig 3e, h**), fasted mice overnight and then provided access to dry chow without water and recorded responses (**Fig 3f, g**). Similar to PVH^AVP^ neurons, we observed a ramping activation of SFO^Glut^ neurons over ∼10 min during eating (**Fig 3i, k**). Examination of the short timescale relationship between individual feeding bouts and SFO^Glut^ activity revealed that young mice consistently showed a spike in SFO activity after feeding bouts terminated, whereas old mice showed no change (**Fig 3i-l**). This difference mirrors the disrupted short timescale coupling between eating and calcium dynamics we observed in PVH^AVP^ neurons (**Fig 2i, j**). Thus, the delayed onset of drinking after feeding in old mice (**Fig 3d**) is consistent with defective SFO activation in response to acute (i.e. presystemic) ingestive signals.

In contrast to these short timescale differences, we observed no difference between young and old animals in the total amount of food consumed (**Fig 4a-c**), the magnitude of SFO^Glut^ neuronal activation over the entire trial (**Extended Data Fig 4a-c, e**), or the time to half maximal SFO^Glut^ activation in males (**Extended Data Fig 4b, d, e**). This indicates that SFO^Glut^ neurons respond similarly to food ingestion in young and old animals on a timescale of ∼30 min. When aged females (but not males) were given reaccess to water (**Fig 4a, b**), we did observe that they drank less than young animals (859 ± 86 vs. 409 ± 56 licks, young vs. aged, P <0.001, **Fig 4d** and **Extended Data Fig 4f**) and we confirmed this result with a second cohort of mice (**Extended Data Fig 4h-j**). Consistent with this behavior, the magnitude of SFO^Glut^ inhibition by water drinking was smaller in aged females compared to young females (-2.87 ± 0.19 vs. -2.30 ± 0.09 z, young vs. aged, P = 0.02, **Fig 4f, g**). However, we are hesitant to ascribe this effect to an age-related decline in thirst, because the difference was driven by unusually high water intake in the young female mice, which exceeded that of young males (859 ± 86 vs. 443 ± 54 licks, female vs. male, P < 0.001, unpaired t-test). This suggests that young female mice show a specific enhancement in prandial thirst, which is lost during aging.

**Figure 4.**
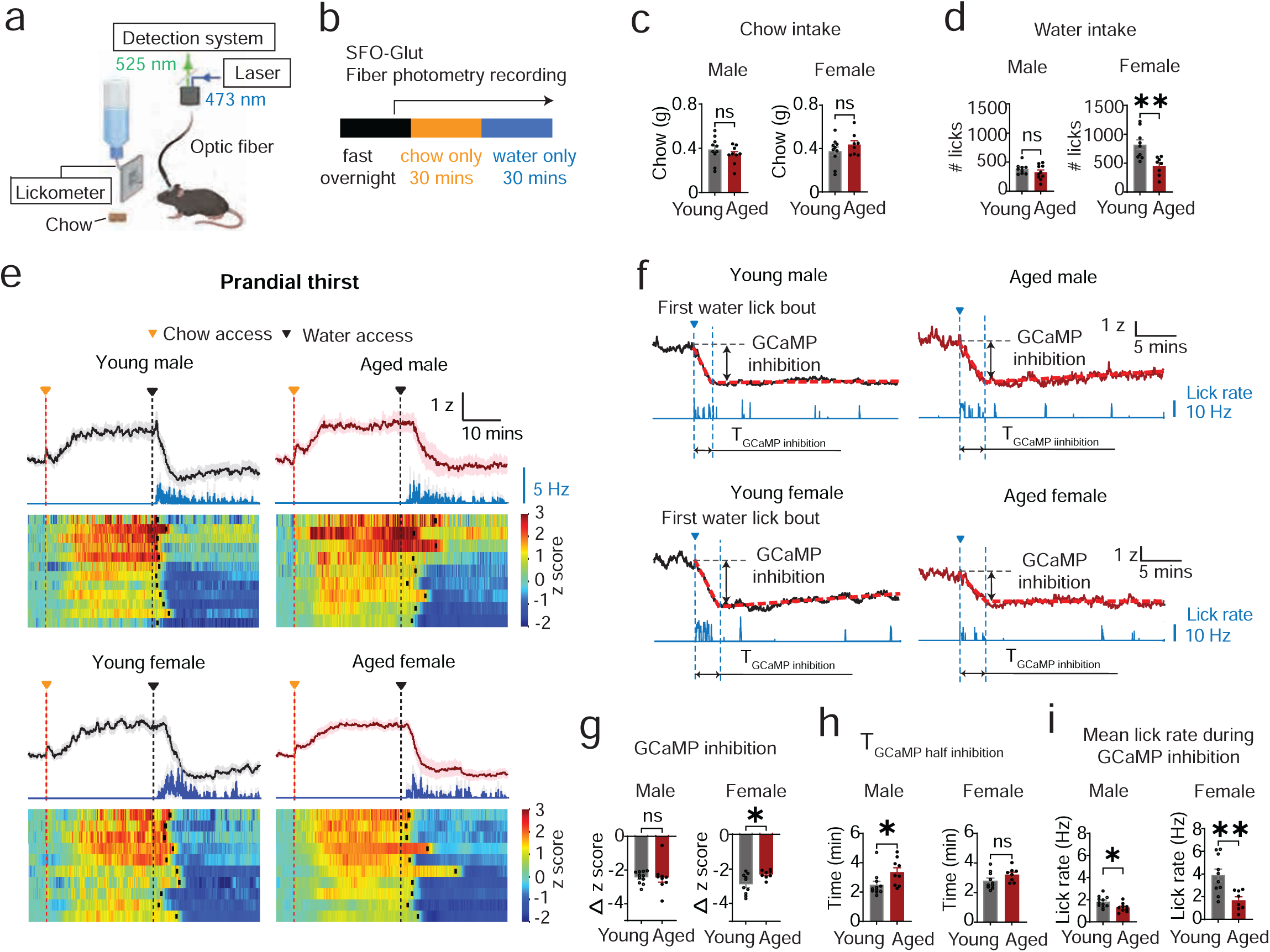
Age reduces water intake after feeding in females and delays SFO^Glut^ neuronal inhibition by rehydration. **a.** Behavioral setup for SFO^Glut^ fiber photometry during dry chow intake and prandial drinking. **b.** Experimental paradigm for SFO^Glut^ fiber photometry during 30 minute chow consumption following overnight fasting, followed by 30 minutes of water access. **c.** Chow intake during 30 minutes of chow access. n.s., non-significant by unpaired t-test. **d.** Total number of water licks during the first 25 minutes following 30 minutes of chow access. n.s., non-significant and **P<0.01 by unpaired t-test. **e.** (Top) Mean lick rate (blue) and mean z-score traces of GCaMP fluorescence in young (2-5 mo, black) and aged (25-27 mo, magenta) male and female mice, aligned by chow access (red dotted vertical line). (Bottom) Heat-maps of individual z-score traces in each group. Black dotted lines represent water access; black rectangle s mark the onset of water lick bout in each recording. **f.** Representative of lick rate (blue) and z-score traces of GCaMP fluorescence during water drinking after feeding, aligned to the first water lick bout (blue dotted line with triangle). Examples are shown from 3- (top left) and 26- (top right) month-old male mice and 4- (bottom left) and 27- (bottom right) month-old female mice. Red dotted lines represent linear regression fits for the fast inhibition phase and steady phase of SFO^Glut^ activity. **g.** Magnitude of SFO^Glut^ inhibition during the fast inhibition phase as indicated in **f.** ns. non-significant and *P<0.05 by unpaired t-test. **h.** Half time for GCaMP to reach the transition from the fast inhibition phase to the steady phase. n.s., non-significant and *P<0.05 by unpaired t-test. **i.** Mean lick rate during the fast SFO^Glut^ inhibition phase. *P<0.05 and **P<0.01 by unpaired t-test.

### Water intake after deprivation increases with age

The preceding data show that aging alters primarily the short timescale coupling between eating and drinking, which is determined by pre-systemic cues from the oropharynx and GI tract^1,2,4,46–48^. To characterize how aging alters responses to systemic signals, we first measured the response of young and old mice to 24-hour water deprivation. We found that water deprivation increased blood osmolality in both young and old mice, and there was no difference in the absolute blood osmolality after deprivation between young and old animals (male: 346 ± 4 vs. 348 ± 5 mOsm/kg, 2-5 mo. vs. 26 mo., P = 0.84; female: 341 ± 4 vs. 347 ± 3 mOsm/kg, 2-5 mo. vs. 26 mo., P = 0.29 by uncorrected Fisher’s LSD, **Fig 5a**). Of note, this was despite the fact that aged mice had a higher starting blood osmolality (**Fig 5a**) and a reduced capacity to concentrate their urine (male: 3500 ± 123 vs. 2485 ± 162 mOsm/kg, 2-5 mo. vs. 26 mo., P < 0.001; female: 3950 ± 257 vs. 2563 ± 122 mOsm/kg, 2-5 mo. vs. 26 mo., P < 0.001 by uncorrected Fisher’s LSD, **Fig 5b** and **Extended Data Fig 5b**). It is possible that these defects in aged mice were compensated for by opposing changes in water loss by other routes, although we observed no differences in chow intake during water deprivation in aged mice compared to young controls (**Extended Data Fig. 5a**).

**Figure 5.**
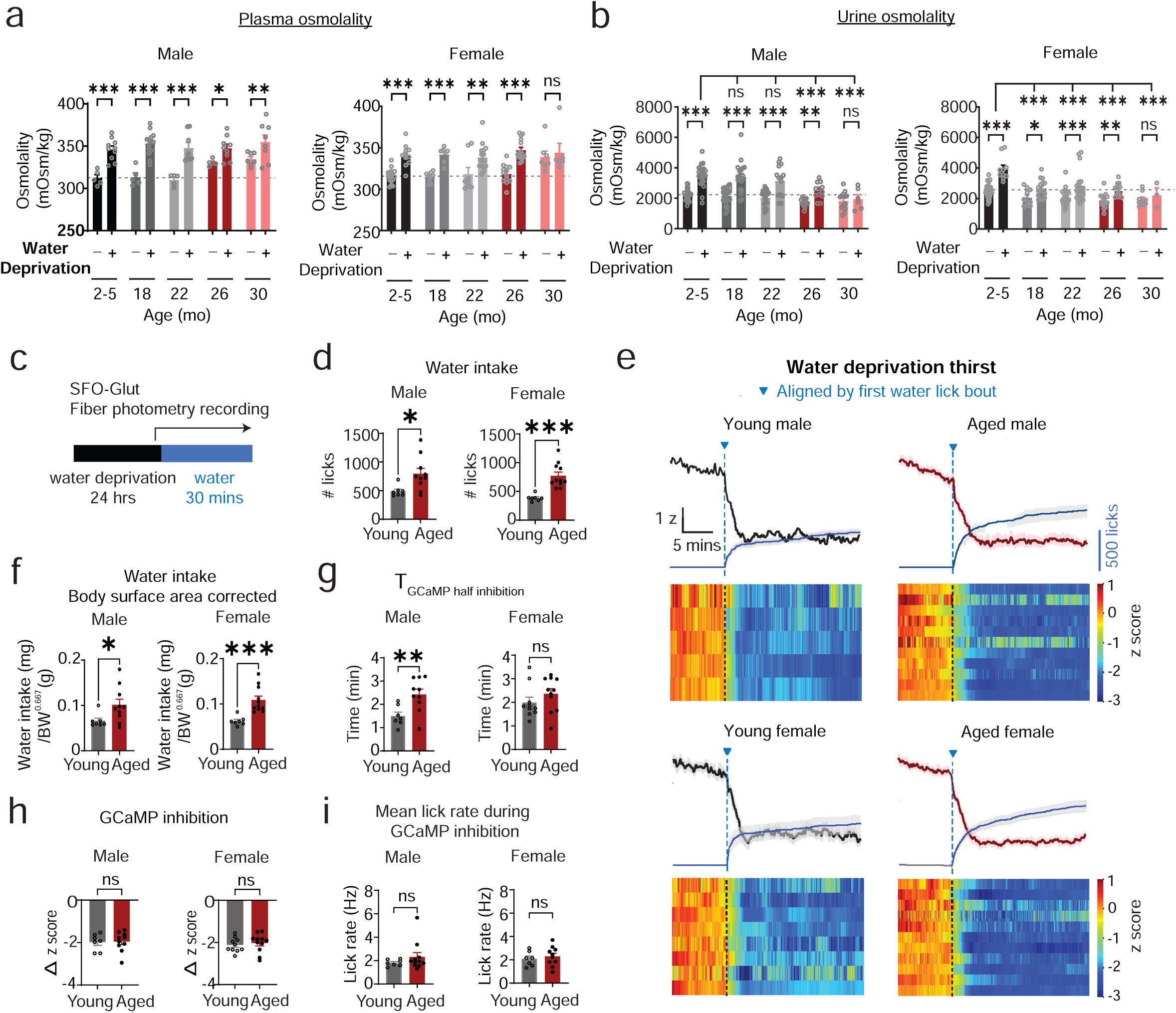
Age increases water intake after deprivation and delays SFO^Glut^ neuronal inhibition by rehydration. **a.** Plasma osmolality under baseline conditions with *ad libitum* access to food and water and after 24 hours of water deprivation in young (2-5 mo) and aged (26-27 mo) male (left) and female (right) mice. ns, non-significant, *P<0.05, **P<0.01 and ***P<0.001 by uncorrected Fisher’s LSD test. For males, ***P<0.001 and *P<0.05 for main effects of hydration state [F(4, 66)=1.886] and age [F(1, 66)=79.26] respectively, and non-significant for their interactions [F(4, 66)=3.403] by 2-way ANOVA. For females, ***P<0.001 and *P<0.05 for main effects of hydration state [F(1, 99)=53.33] and age [F(4, 99)=2.493] respectively, and non-significant for their interactions [F(4, 99)=1.715] by 2-way ANOVA. **b.** Urine osmolality during under baseline conditions and after 24 hours of water deprivation in young (2-5 mo) and aged (26-27 mo) male (left) and female (right) mice. ns, non-significant, *P<0.05, **P<0.01 and ***P<0.001 by Fisher’s LSD test. For males, ***P<0.001 for main effects of hydration state [F(4, 161)=69.12] and age [F(4, 161)=10.85], and *P<0.05 for interactions of their main effects [F(4, 161)=3.056] by 2-way ANOVA. For females, ***P<0.001 for main effects of hydration state [F(1, 135)=37.47] and age [F(4, 135)=12.88], and *P<0.05 for their interactions [F(4, 135)=2.670] by 2-way ANOVA. **c.** Behavioral paradigm for SFO^Glut^ fiber photometry during 30 minutes of water intake following 24 hours of deprivation. **d.** Total number of water licks during 25 minutes. *P<0.05 and ***P<0.001 by unpaired t-test. **e.** (Top) Mean cumulative number of licks (blue) and mean z-score trace of GCaMP fluorescence in young (3-6 mo, black) and aged (26-28 mo, magenta) male and female mice, aligned to the first water lick bout (blue dotted line). (Bottom) Heatmaps of individual z-score traces. **f.** Total water intake during 25 minutes, normalized to the body surface area (BW^0.667^). *P<0.05 and ***P<0.001 by unpaired t-test. **g.** Half time for GCaMP signals to reach the transition from the fast SFO^Glut^ inhibition phase to the steady phase. n.s., non-significant and **P<0.01 by unpaired t-test. **h.** Magnitude of SFO^Glut^ inhibition during the fast inhibition phase as indicated in **e.** n.s., non-sifnigicant by unpaired t-test. **i.** Mean lick rate during the fast SFO^Glut^ inhibition phase. n.s., non-significant by unpaired t-test.

We next measured water intake when mice were given re-access to water for 30 min following 24-hour deprivation (**Fig 5c**). Surprisingly, we found that aged male and female mice consumed more water than their young counterparts in this test (male: 489 ± 36 vs. 795 ± 93 licks, young vs. aged, P = 0.01; female: 463 ± 66 vs. 722 ± 46 licks, young vs. aged, P = 0.005, **Fig 5d, e**). This difference persisted when we normalized the water intake to the body surface area (male: 66.2 ± 5.8 vs. 101.4 ± 12.5 mg/g^0.667^, young vs. aged, P = 0.04; female: 62.3 ± 3.2 vs. 108.9 ± 8.6 mg/g^0.667^, young vs. aged, P < 0.001, **Fig 5f**), which has been proposed as a method to correct for differences between animals in body size^19,20^. Thus, although the plasma osmolality of young and aged mice is similar after 24-hour deprivation, aged mice drink more when water is again made available.

We considered two hypotheses, which are not mutually exclusive, for why aged mice would drink more after water deprivation. The first possibility is that there is a defect in thirst satiation in old animals. Thirst satiation is encoded in the inhibition of SFO^Glut^ neurons, which show stepwise decreases in activity during drinking that track the volume of fluid consumed^18^. If aged mice have a defect in thirst satiation, then we would expect SFO^Glut^ neurons to be inhibited more slowly when aged mice drink. Indeed, we observed that SFO^Glut^ neurons showed a delayed inhibition when aged male mice drank after 24-hour deprivation (time to half inhibition: 1.5 ± 0.2 vs. 2.4 ± 0.2 min, young vs. aged, P = 0.008, **Fig 5g**), and a similar trend was observed in aged females (2.0 ± 0.2 vs. 2.4 ± 0.2 min, young vs. aged, P = 0.26, **Fig 5g**). Importantly, this difference was not due to differences in behavior, because young and aged mice drank at the same rate when water was made first available (**Fig 5i**). This suggests that the overdrinking of old mice may be caused, in part, by an age-related impairment in the presystemic sensory feedback that inhibits SFO^Glut^ neurons during water consumption. Consistent with this result, we found that the inhibition of PVH^AVP^ neurons by water drinking after 24 hour deprivation was also delayed in aged mice compared to young controls (1.14 ± 0.06 vs 1.82 ± 0.55 min, young vs aged, P = 0.03, **Extended Data Fig 6a-d**), despite comparable drinking rates (2.7 ± 0.4 vs 2.7 ± 0.6 Hz, young vs aged, P = 0.99, **Extended Data Fig 6e**).

### Functional manipulations suggest greater SFO activation in aged animals after deprivation

An alternative, and not mutually exclusive, explanation for why aged animals drink more after water deprivation is that the thirst circuitry is activated more strongly. In this regard, while photometry can measure relative changes in response to a stimulus, it cannot reliably quantify absolute differences in activity between cohorts of animals. Thus, it is possible that aged animals, despite similar blood osmolality as young animals, have greater activation of the thirst circuitry after water deprivation and this causes them to drink more.

We performed three sets of experiments to test this hypothesis. First, we took advantage of the fact that optogenetic stimulation of SFO^Glut^ neurons creates a state of virtual thirst that drives drinking in water-sated animals^17,18,21,49,50^. Therefore, by optogenetic stimulation of SFO^Glut^ neurons, we can introduce a fixed amount of activation into the thirst circuit of young and old animals and then compare the behavioral responses. We targeted ChR2 to SFO^Glut^ neurons of young and old male mice by viral injection of a Cre-dependent AAV and implanted an optical fiber above the SFO (**Fig 6a**). We first water-deprived these animals for 24 hours and then measured intake for 30 min in the absence of laser stimulation. Aged mice consumed more water than young animals after 24-hour deprivation (426 ± 43 vs. 748 ± 163 licks, young vs. aged, P = 0.03, **Fig 6c**), mirroring our findings from a different cohort of mice used for photometry experiments (**Fig 5d**). We then optogenetically stimulated the same animals in the water-sated state and measured drinking (**Fig 6b**). Strikingly, we found that there was no difference in the amount of water consumed between young and old animals after artificial activation of the SFO^Glut^ (762 ± 136 vs. 757 ± 245 licks, young vs. aged, P = 0.98, unpaired t-test, **Fig. 6d**). The fact that we observe no difference in this test is consistent with the hypothesis that the greater water consumption in aged animals is due to greater activation of this circuitry following 24-hour deprivation.

**Figure 6.**
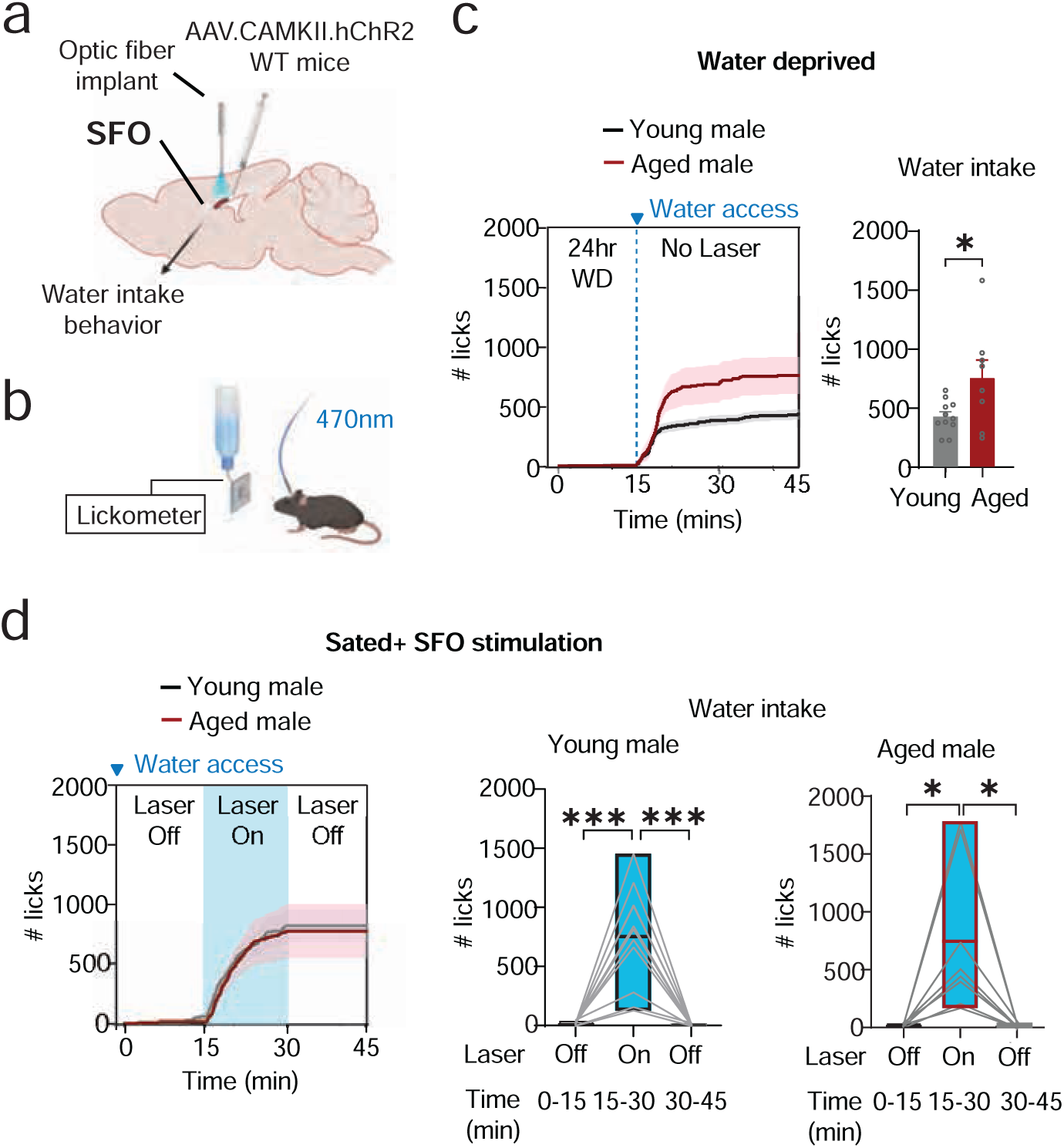
Age preserves the SFO^Glut^ synaptic output for water intake behavior. **a.** Schematic illustrating stereotaxic injection of AAVs expressing channelrhodopsin-2 (ChR2) into the SFO and implantation of an optic fiber for optogenetic stimulation. **b.** Schematic of optogenetic stimulation of SFO^Glut^ neurons and recording of water intake behavior. **c.** (Left) Cumulative number of licks in mice subjected to 24 hours of water deprivation. Water access was provided between 15-45 minutes of behavioral recording and no laser stimulation was delivered. Total number of licks during the first 30 minutes of water access in young (2-7 mo, middle) and aged (26-29 mo, right) male mice. P<0.05 by unpaired t-test. **d.** (Left) Cumulative number of licks in water-sated mice. Blue light stimulation (470 nm, 20 Hz) was applied between 15-30 minutes (blue shaded area). (Middle, right) Total number of licks in young and aged male mice during laser-off (0-15 min), laser-on (15-30 min), and post-stimulation (30-45 min) periods. *P<0.05 and P<0.001 by paired t-test.

To test this hypothesis a second way, we asked whether there are differences between young and old animals in their motivation to work for water. We reasoned that if aged mice drink more because they have greater SFO activation, then they should also lever press more in an operant task for water, since this is controlled by SFO activity^50,51^.

We trained mice on a fixed-ratio 1 (FR1) schedule, so that they received one drop of water for each press of the active lever but not the inactive lever (**Fig. 7a**). We then water-deprived animals for 24 hours and measured lever pressing in a 30 min test. Strikingly, we found that aged male and female mice both engaged in more lever pressing for water compared to young controls (male: 55 ± 6 vs. 89 ± 9, young vs. aged, P = 0.02, female: 49 ± 5 vs. 73 ± 7, young vs. aged, P = 0.04, **Fig. 7b, c**). This enhanced responding was specific to the active lever, confirming that old mice have heightened motivation specifically for water. Importantly, both young and old mice consumed significantly less water during the FR1 test compared to free drinking (**Extended Data Fig 7a**), implying that differences in satiation are unlikely to explain this difference in lever pressing. To exclude any contribution from differences in satiation, we performed a progressive ratio test (PR3) in which animals were required to perform an increasing number of lever presses for each subsequent water reward, such that they reach a “break point” long before their thirst has been significantly reduced. In this test, we observed again that old male mice lever pressed more vigorously than young controls (break point: 8 ± 1 vs. 15 ± 2, young vs. aged, P = 0.02, **Fig 7d**). In females, there was a trend to greater lever pressing that was non-significant (8 ± 1 vs. 10 ± 1, young vs. aged, P = 0.33, **Fig 7d**). However, we acknowledge that the increased lever pressing in aged mice could still be partly explained by age-related differences in the rate of satiation, whereby each individual reward may produce weaker inhibition of SFO^Glut^ neuronal activity in aged animals, allowing these neurons to remain relatively more active throughout the task. Even with this caveat, the combined FR1 and PR3 data support the notion that aged mice show increased motivation for water following 24-hour deprivation, consistent with stronger activation of thirst circuitry in this context.

**Figure 7.**
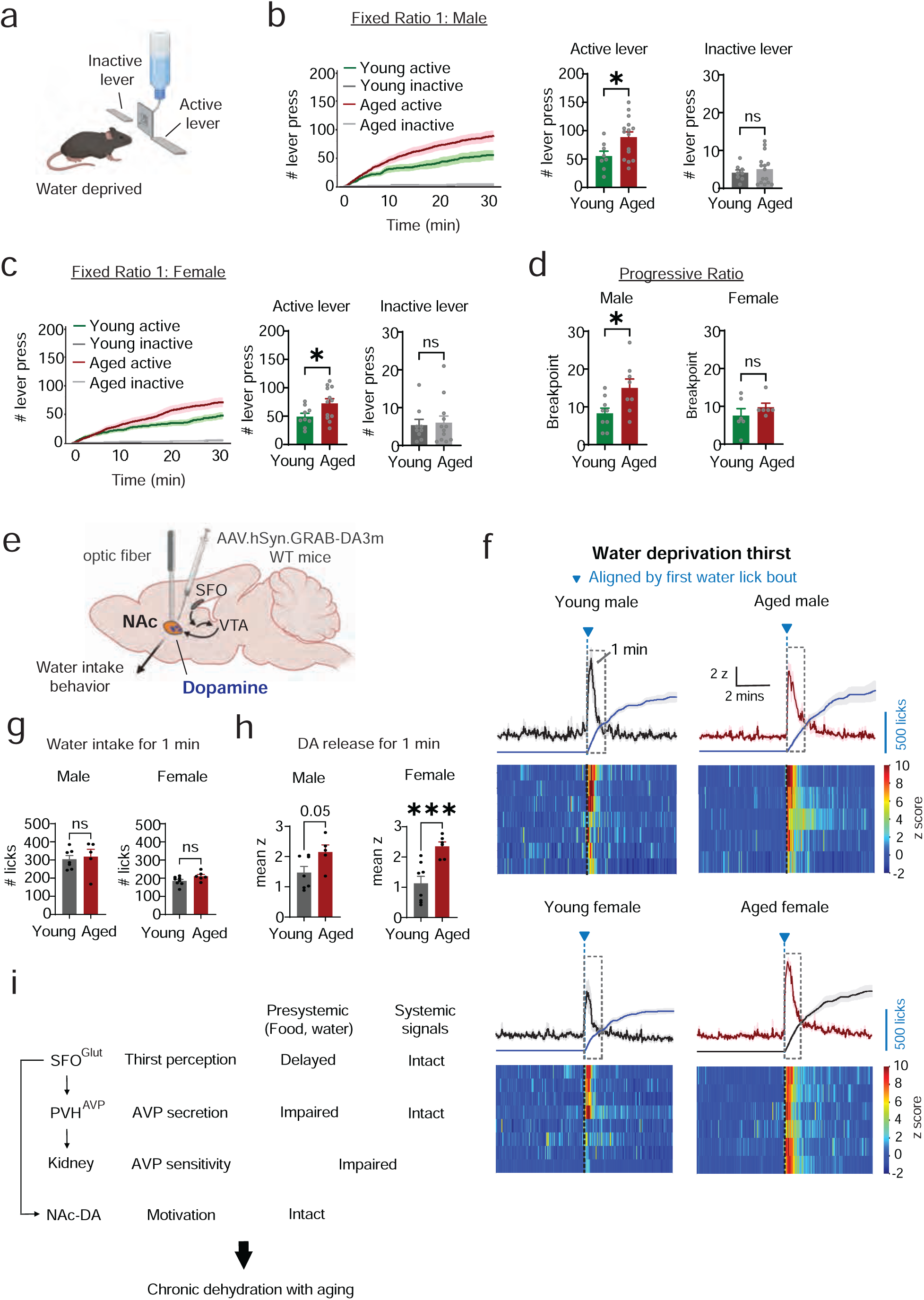
Age increases motivation for water after deprivation. **a.** Schematic of the lever-press operant task following 24 hours of water deprivation. **b.** Fixed Ratio 1 (FR1) in operant task for water for 30 minutes in water-deprived young (2-5 mo, green) and aged (23-28 mo, magenta) male mice. (Left) Cumulative number of lever presses over time. (Middle) Total number of active lever presses in 30 minutes. (Right) Total number of inactive lever presses in 30 minutes. ns, non-significant and *P<0.05 by unpaired t-test. **c.** Fixed Ratio 1 (FR1) in operant task for water for 30 minutes in water-deprived young (2-6 mo, green) and aged (23-27 mo, magenta) female mice. (Left) Cumulative number of lever presses over time. (Middle) Total number of active lever presses in 30 minutes. (Right) Total number of inactive lever presses in 30 minutes. ns, non-significant and *P<0.05 by unpaired t-test. **d.** Breakpoint values from a Progressive Ratio 3 (PR3) operant task for water in water-deprived young and aged male and female mice. ns, non-significant and *P<0.05 by unpaired t-test. **e.** Schematic illustrating stereotaxic injection of the GRAB-DA dopamine sensor into the nucleus accumbens (NAc) and implantation of an optic fiber for fiber photometry. SFO, subfornical organ. VTA, ventral tegmental area. Arrows denote synaptic projections. **f.** GRAB-DA fiber photometry during water intake following 24 hours of water deprivation. (Top) Cumulative number of licks (blue) and mean z-score trace of GRAB-DA fluorescence in young (3-6 mo, black) and aged (24-26 mo, magenta) male and female mice, aligned to the first water lick bout (blue dotted line). (Bottom) Heatmaps of individual z-score traces, aligned to the first water lick bout (black dotted line). **g.** Number of water licks during the first water lick bout shown in **f.** *P<0.05 by unpaired t-test. **h.** Area under the curve of z-score of GRAB-DA fluorescence during the first 3 minutes following the initial water lick bout after 24-hour water deprivation shown in **f.** **P<0.01 by unpaired t-test. **i.** Schematic summary of age-related neural activity changes observed across thirst circuit recordings.

Motivation for water is controlled, in part, by dopamine release in the nucleus accumbens (NAc). Cues that predict water, including its taste, trigger a burst of DA in NAc that helps to orient and energize further actions toward obtaining water^52–54^. The magnitude of this DA release is controlled by SFO^Glut^ neuron activity and is greater in thirsty animals^53,54^. We therefore measured water-triggered DA release in NAc as an independent measure of whether old animals have greater activation of the thirst circuitry following 24-hour deprivation.

We targeted the genetically encoded DA sensor GRAB-DA3m to the NAc and, in the same surgery, implanted an optical fiber above the NAc to record DA release by fiber photometry (**Fig. 7e**). Mice were water deprived for 24 hours and then given access to a water bottle for self-paced drinking. The onset of drinking was accompanied by a spike in DA release in both young and old animals, but this increase was bigger (**Extended Data Fig 7c**) and lasted longer (time to half decline, male: 38.1 ± 3.3 vs. 74.8 ± 17.4 s, P = 0.03; female: 35.2 ± 2.8 vs. 66.2 ± 10.4 s, P = 0.007, **Extended Data Fig 7d**) in aged mice. In contrast, there was no difference in the DA response when young and old mice were presented with an empty water bottle, indicating that this effect is specific to water (**Extended Data Fig 7e, f**). Of note, aged mice drank more water during the 30 min test after deprivation (**Extended Data Fig 7b**), as observed previously (**Fig 5d**, **6c**), but differential DA release was observed even at early time points (i.e. 1 minute from the onset of the first drinking bout: male: 1.47 ± 0.20 vs. 2.15 ± 0.24 z, young vs. aged, P = 0.05; female: 1.13 ± 0.23 vs. 2.35 ± 0.15 z, young vs. aged, P < 0.001, **Fig 7f, h**) when the amount drunk was similar (male: 304 ± 20 vs. 318 ± 41 licks, young vs. aged, P = 0.76; female: 185 ± 9 vs. 211 ± 10 licks, young vs. aged, P = 0.08, **Fig 7g**), indicating that this cannot be attributed to differences in behavior. Taken together, these results show that, following 24-hour water deprivation, aged mice drink more (**Fig 5d** and **Extended Data Fig 7b**), show greater motivation to work for water (**Fig 7b-d**), and show greater water-triggered DA release in NAc (**Fig 7h**). This supports the idea that the core thirst circuitry is more strongly activated following 24-hour deprivation in aged mice.

### Aged mice have intact water consumption and delayed SFO inhibition after osmotic thirst

Water deprivation results in increased blood osmolality and decreased blood volume (**Fig 5a** and **Extended Fig 8c**), both of which promote thirst. To characterize how these two pathways are altered by age, we independently manipulated them and measured drinking behavior and SFO^Glut^ neuronal responses.

We first investigated the role of blood osmolality by challenging mice with salt (intraperitoneal injection of 2M NaCl), which we confirmed increases plasma osmolality without significantly altering blood volume^55^ (**Extended Data Fig 8a**). Salt challenge activated SFO^Glut^ neurons in young and old mice to a similar degree (**Fig 8c, Extended Data Fig 8d, f**) and with similar kinetics (**Extended Data Fig 8e**). We then made water available and recorded responses during 30 min of self-paced drinking (**Fig 8a, d-g**). In contrast to 24-hour deprivation, we observed no significant age-related differences in water intake in either male (756 ± 93 vs. 784 ± 135 licks, young vs. aged, P = 0.8, **Fig 8b**) or female mice (875 ± 94 vs. 1065 ± 185 licks young vs. aged, P = 0.33, **Fig 8b**). We replicated this finding in a separate cohort of young and old mice (**Extended Data Fig 8g**). However, when we examined SFO^Glut^ neuronal recordings, we found that the activity of these neurons during drinking declined more slowly in aged animals (male: 2.4 ± 0.4 vs. 6.4 ± 1.1 min, young vs. aged, P = 0.009; female: 1.8 ± 0.2 vs. 4.4 ± 1.0 min young vs. aged, P = 0.02, **Fig 8e**). This was not due to differences in lick rate in young and old animals (**Fig 8f**) and mirrors the finding of slower SFO^Glut^ inhibition when old animals drink after 24-hour deprivation (**Fig 5g**). Together, these data indicate that, in aged animals, SFO^Glut^ neurons can sense changes in blood osmolality appropriately, but that there is a delay in their inhibition by presystemic cues during drinking.

**Figure 8.**
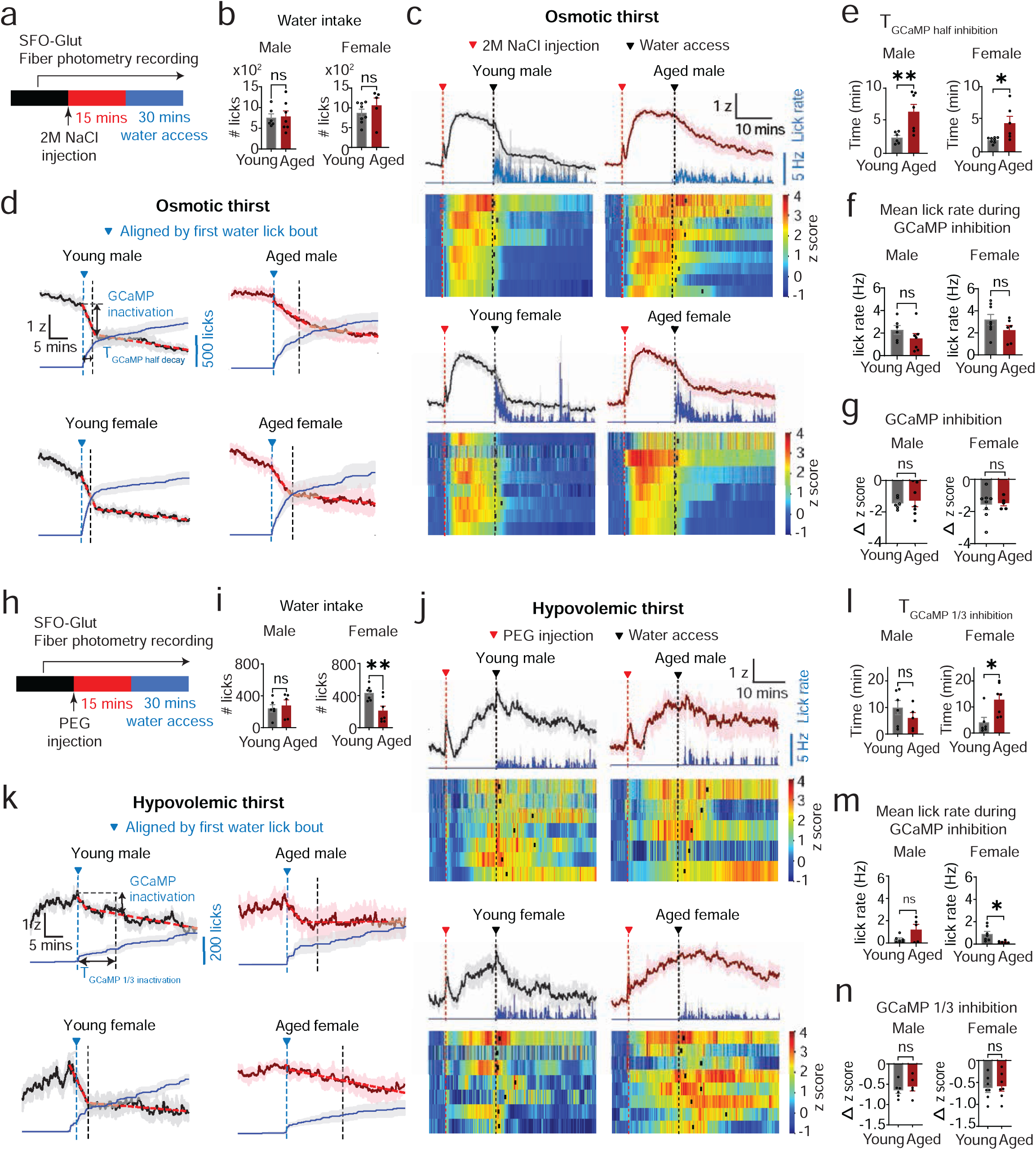
Age delays SFO^Glut^ neuronal inhibition by rehydration after osmotic and hypovolemic thirst. **a.** Experimental paradigm for SFO^Glut^ fiber photometry and behavioral recording for osmotic thirst induced by acute intraperitoneal 2 M NaCl injection. **b.** Total number of water licks during 25 minutes. n.s., non-significant. **c.** (Top) Mean lick rate (blue) and mean z-score trace of GCaMP fluorescence in young (4-7 mo, black) and aged (24-29 mo, magenta) male and female mice, aligned to 2 M NaCl injection (red dotted line). (Bottom) Heatmaps of individual z-score traces. Black dotted lines indicate water access, and black rectangles mark the onset of the first drinking bout. **d.** Cumulative number of water licks (gray) and mean z-score of GCaMP fluorescence in young (black) and aged (magenta) male and female mice, aligned to the first water lick bout (blue dotted line with triangle) **e.** Time for GCaMP signals to decay from peak to its half value of z-score measured between the first lick (t = 0; blue dotted line) and the subsequent steady phase (black dotted line) as indicated in **d**. *P<0.05 and **P<0.01 by unpaired t-test. **f.** Mean lick rate during the fast SFO inhibition phase. n.s., non-significant by unpaired t-test. **g.** Magnitude of SFO inhibition during the fast SFO inhibition phase as indicated in **d.** n.s., non-significant by unpaired t-test. **h.** Experimental paradigm for SFO^Glut^ fiber photometry recording for hypovolemic thirst induced by acute subcutaneous PEG injection. **i.** Total number of water licks during 25 minutes. n.s., non-significant and **P<0.01 by unpaired t-test. **j.** (Top) Mean lick rate (blue) and mean z-score trace of GCaMP fluorescence in young (4-7 mo, black) and aged (25-29 mo, magenta) male and female mice, aligned to PEG injection (red dotted line). (Bottom) Heatmaps of individual z-score traces. Black dotted lines indicate water access, and black rectangles mark the onset of the first drinking bout. **k.** Cumulative number of water licks (gray) and mean z-score of GCaMP fluorescence in young (black) and aged (magenta) male and female mice, aligned to the first water lick bout (blue dotted line with triangle). **l.** Time for GCaMP signals to decay from peak to its 1/3 value of z-score measured between the first lick (t = 0; blue dotted line) and the subsequent steady phase (black dotted line) as indicated in **k**. n.s., non-significant and *P<0.05 by unpaired t-test. **n.** Magnitude of SFO inhibition during decay from peak to its 1/3 value of peak fluorescence as indicated in **k.** n.s., non-significant.

### Water consumption after hypovolemia is selectively reduced in aged females

We next investigated how aging affects responses to decreases in blood volume. To do this, we used a paradigm^56^ in which mice are given subcutaneous injection of polyethylene glycol (PEG), which forms an edema under the skin that draws water out of the circulation and thereby reduces blood volume. We found that PEG injection resulted in similar decrease in blood volume in young and old mice (**Extended Data Fig 8b**). In SFO^Glut^ photometry recording, PEG injection caused a ramping activation of SFO^Glut^ neurons, and there was no difference in the magnitude or rate of this activation in young versus old mice (**Fig 8j**; **Extended Data Fig 8h-j**). When mice were subsequently given reaccess to water for 30 min (**Fig 8h, k-n**), we found old females, but not males, consumed much less water than their young counterparts (**Fig 8i**), which we confirmed in an independent cohort of young and old mice (**Extended Data Fig 8k**). Interestingly, these differences between young and old female mice were driven by unusually high intake in the young females, which exceeded that in young males (433 ± 34 vs. 241 ± 47 licks, female vs. male, P = 0.006, unpaired t-test, **Fig 8i**). This mirrors what we observed in assays of prandial thirst (**Fig 4d**), in which young females consumed much more water than other cohorts, and suggests that food intake and PEG-induced hypovolemia may act through a common mechanism to cause overdrinking in young females. One candidate is the hormone Angiotensin II (AngII), which is implicated in both prandial drinking and hypovolemia^21,57–59^, but we observed no difference between young and old animals in the SFO^Glu^ response to AngII injection (**Extended Data Fig 4k, l**).

## DISCUSSION

Deterioration of fluid homeostasis is one of the hallmarks of aging, but the underlying causes are not well understood. In this study, we have systematically investigated how aging alters fluid balance in the mouse, comparing young and old animals of both sexes across a range of physiologic, behavioral, and neural circuit analyses. This has revealed that aging induces a complex, but reproducible, constellation of fluid homeostasis phenotypes in the mouse. These include (1) chronic dehydration that begins at 26-30 months of age, (2) decreased water resorption by the kidney, secondary to both AVP deficiency and resistance, (3) impaired short-timescale coupling between eating and drinking, both behaviorally and at the level of neural dynamics, (4) delayed inhibition of thirst circuits during drinking, and, surprisingly, (5) an increase in water intake, as well as thirst motivation, in aged animals after 24-hour water deprivation, but not in response to any other dipsogenic stimuli.

### Aged mice have a primary defect in AVP signaling

We found that aged mice of both sexes exhibited chronic dehydration, and this was accompanied by consistent deficits in AVP secretion and function. These deficits included (1) dilute urine, indicating a reduced ability of the kidney to concentrate urine, (2) reduced AVP levels, despite blood hyperosmolality, and (3) reduced sensitivity to supraphysiologic AVPR stimulation, indicating AVP resistance. Because the defects in fluid consumption we observed were mostly acute in nature (discussed below), this suggests that the dehydration in aged mice is driven primarily by deterioration of the AVP-kidney axis. Age-related declines in renal AVP sensitivity are thought to arise from reductions in V2 receptor signaling efficiency, leading to impaired AVP-mediated cAMP/PKA activation and decreased expression of AVP-regulated aquaporins and solute transporters, ultimately compromising the kidney’s urinary concentrating capacity^35,60–64^.

Previous studies have reported mixed findings on the effect of aging on AVP function in different species. Most studies have been performed in rats, where it has been reported that old animals have fewer AVP cells in the PVH and reduced expression of AVP^65,66^. While some studies report reduced circulating levels of AVP^65^, others have found old animals have normal or even enhanced levels of AVP in the blood^66–72^. Similar variability has been found in humans^7,73–76^. These differences could have many causes, including the age of the animals (we used 26-30 month-old mice, which are older than the animals used in many studies); the species, strain, and sex; and the diet on which the animals are maintained over the lifetime. Although we cannot reconcile all these differences, our findings establish a baseline for how aging effects AVP and kidney function in the C57BL/6 mouse strain, which is a commonly used model organism for aging research.

To probe the source of age-related changes in AVP secretion, we recorded calcium dynamics in PVH^AVP^ neurons. This PVH population includes both magnocellular neurons, which release AVP into the blood, and parvocellular neurons that release AVP within the brain. Because we tested these neurons with hydrational stimuli that strongly activate magnocellular cells, we believe that most of the responses we observe reflect magnocellular activity. However, we cannot exclude some parvocellular contribution to the recorded signals. Although in principle this problem could be addressed by recording from AVP neurons in the SON, which are exclusively magnocellular, we were unable to reliably target the SON in very old animals, possibly due to its small size and deep brain location combined with changes in brain anatomy that occur with aging.

### Aging primarily impairs responses to presystemic thirst signals

The signals that control thirst are traditionally divided into two classes: (1) systemic signals that arise from the blood (i.e. hyperosmolality and hypovolemia,) and which are primarily involved in generating thirst after water deprivation^1,3,4^ and (2) presystemic signals that arise from the oropharynx and gastrointestinal (GI) tract which are primarily involved in the rapid control of thirst by ingestion (i.e. the quenching of thirst during drinking and the stimulation of thirst during eating)^1,2,4,46,47^. A basic answered question regards how these two classes of signals are differentially affected by aging.

We examined the response of young and old mice to presystemic and systemic thirst signals at the level of both behavior and neural circuit dynamics. The general finding from these experiments was that aged mice show evidence of impaired responses to ingestion (eating and drinking) but not to manipulations of the blood (summarized in **Fig 7i**). For example, aged mice had a consistent delay in the short timescale coupling between eating food and subsequent water intake, and this delay was accompanied by altered calcium dynamics during eating for both PVH^AVP^ and SFO^Glut^ neurons (**Fig 2e**, **3j**, **3l**). There was also an impairment in the rapid inhibition of PVH^AVP^ and SFO^Glut^ neurons during drinking, such that this inhibition was slower (**Fig 4h**, **5g; Extended Data Fig 2f, 6d**) and less efficient (i.e. reduced Δ z per lick; **Extended Data Fig 4g**). In some experiments, these impairments may have contributed to overdrinking in aged animals (**Fig 5d**). The magnitude of these acute defects was moderate, and in most cases aged animals were able to compensate by adjusting behavior on longer timescales (e.g. **Extended Data Fig 3a**). Nonetheless, we observed these changes across multiple cohorts of aged mice of both sexes, suggesting that they are a general feature of aging of the fluid homeostasis system. Of note, because presystemic signals are detected by sensory innervation of the mouth, throat, and GI tract, these changes could reflect the age-dependent deterioration of these primary sensory afferents. This would mirror the well-known age-dependent changes in primary sensory neurons in other sensory systems, such as vision and hearing^77,78^.

In contrast, we saw little difference between young and old animals following direct manipulations of the blood. Thus, young and old mice of both sexes responded normally to increases in blood hyperosmolality, both in terms of the amount of water consumed (**Fig 8b**) and the response to SFO^Glut^ and PVH^AVP^ neurons (**Fig 8c-g, Extended Data Fig 6g**). For hypovolemia, we observed normal responses in aged male mice, whereas aged females consistently consumed less water than young animals (**Fig 8i**). However, as noted in the text, we are hesitant to ascribe this to an age-dependent defect, because the difference arose from exaggerated fluid intake in the young females when compared to males of the same age. One possible explanation for this observation is that it reflects an adaption of the fluid homeostasis system to the demands of reproduction, which results in an increase in bodily fluid volume of 45% during pregnancy^79–81^. This may make female mice of reproductive-age hypersensitive to signals of hypovolemia. Sex differences from our study are described in Table 1.

**Table 1.**
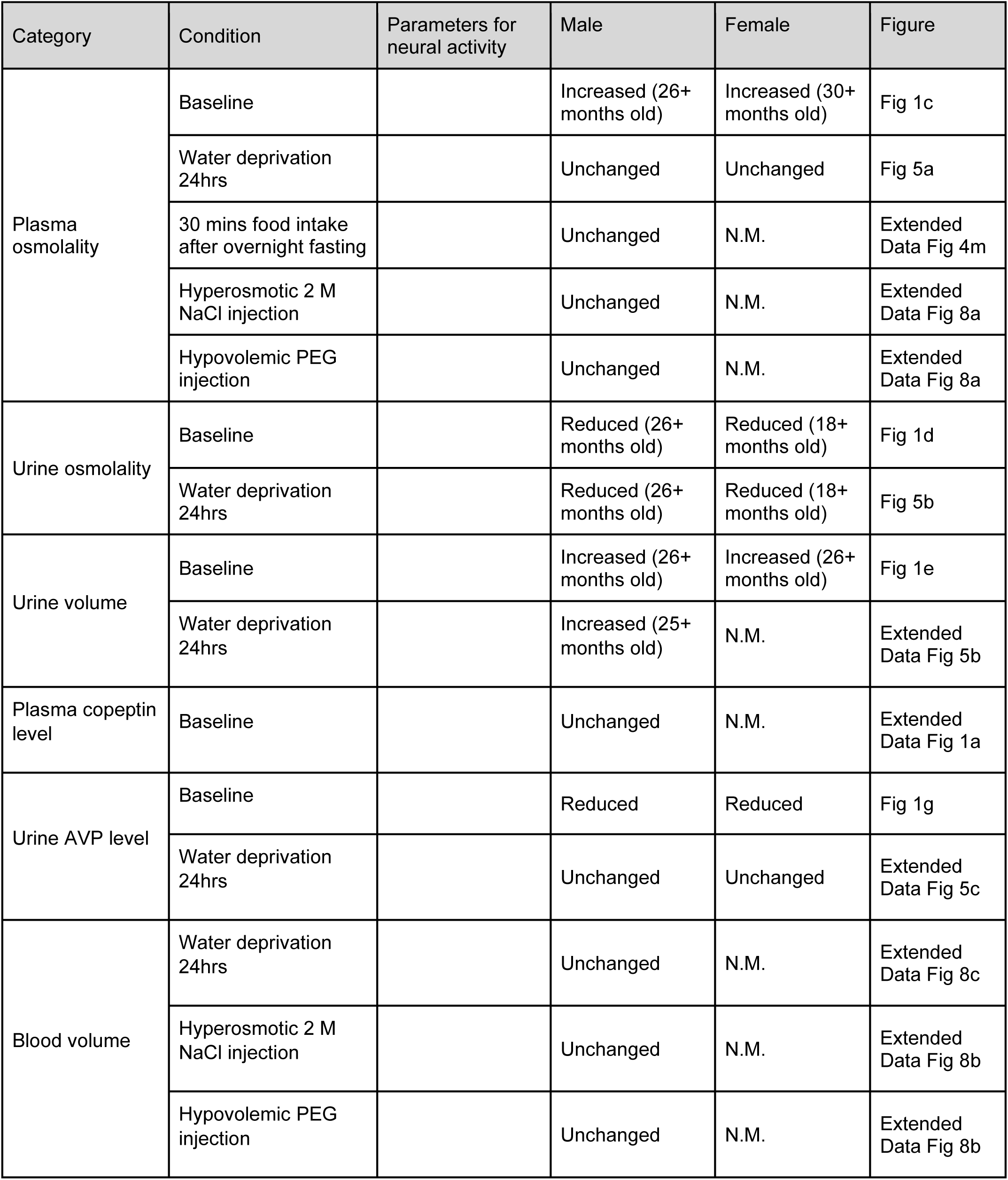
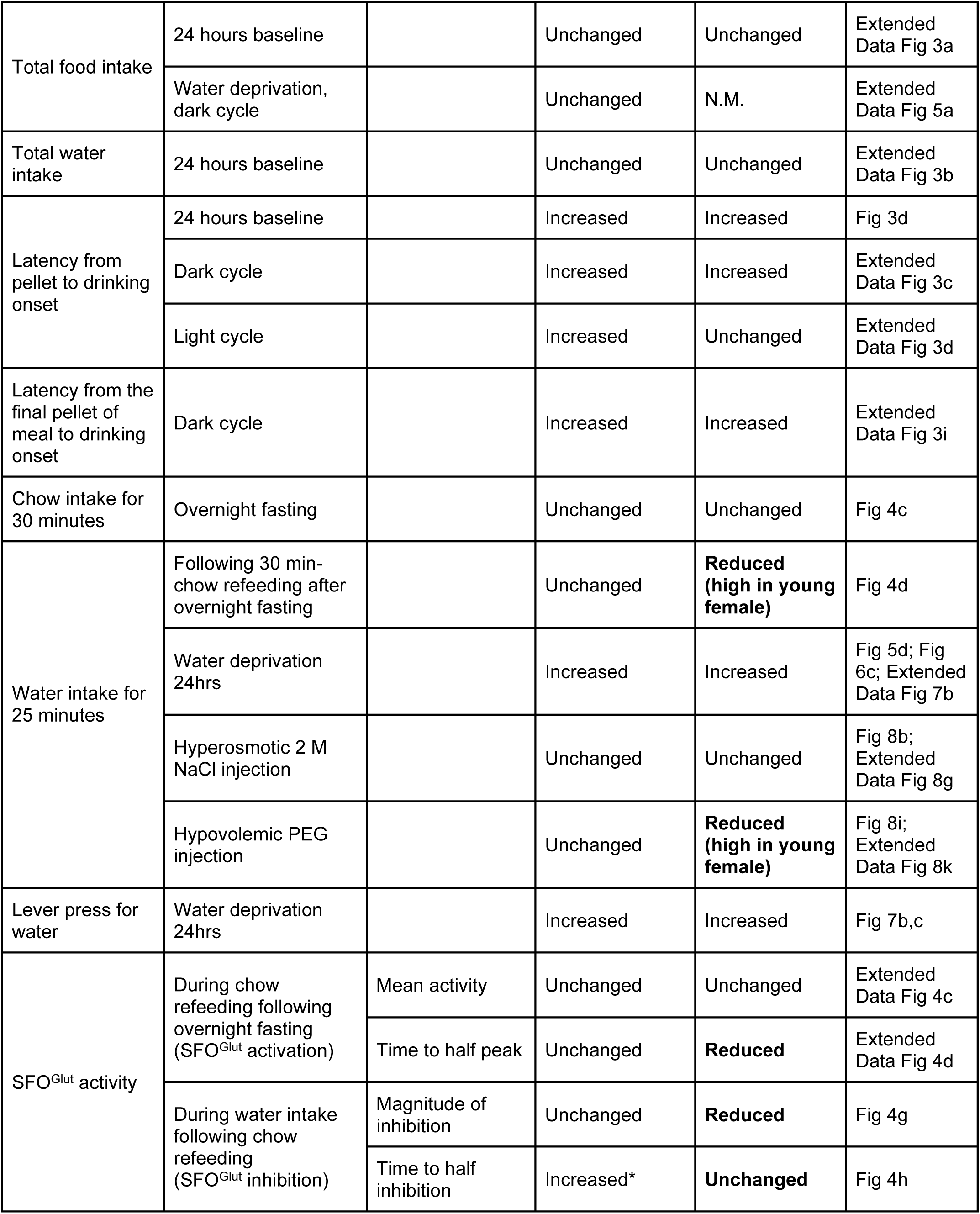
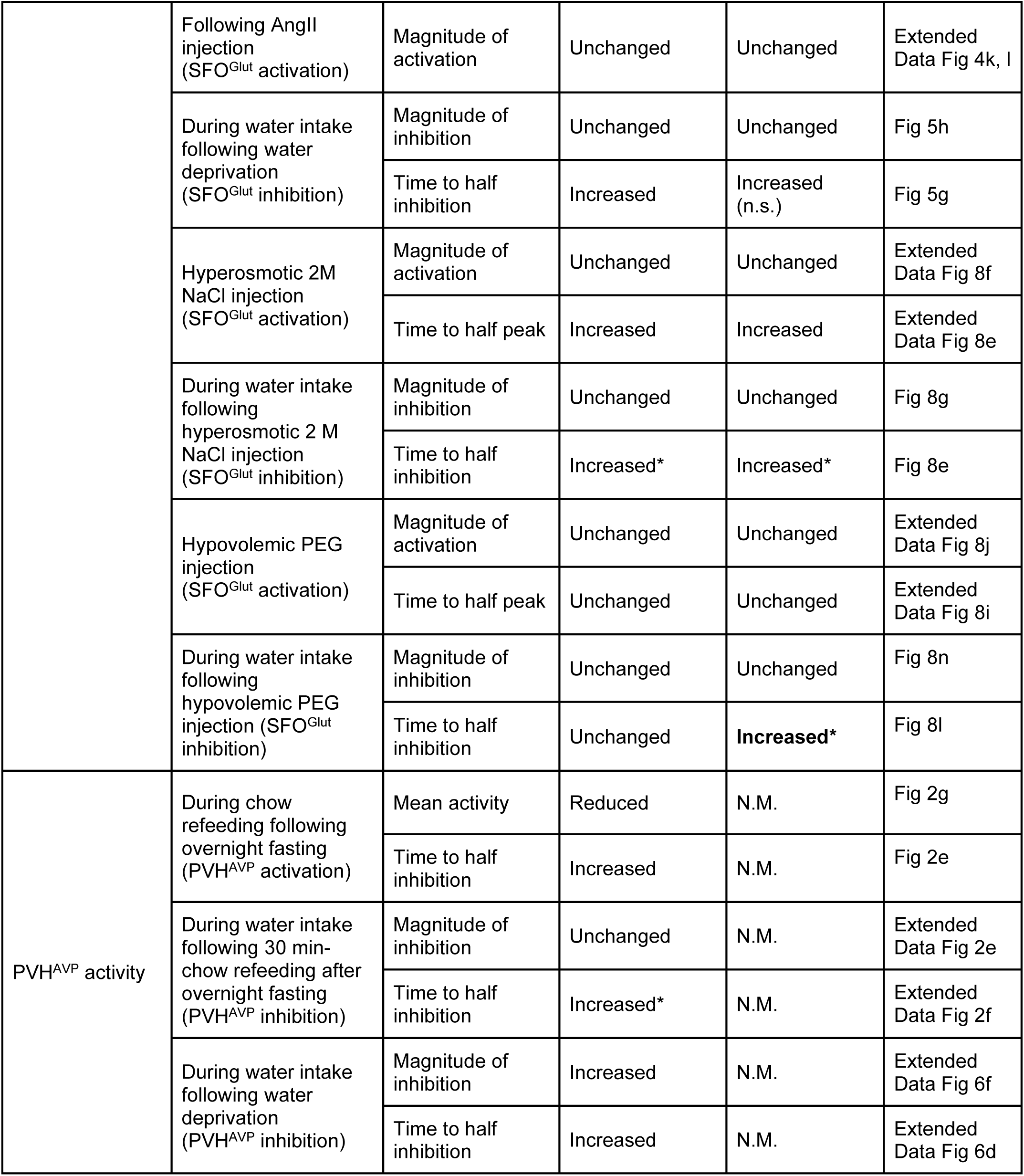
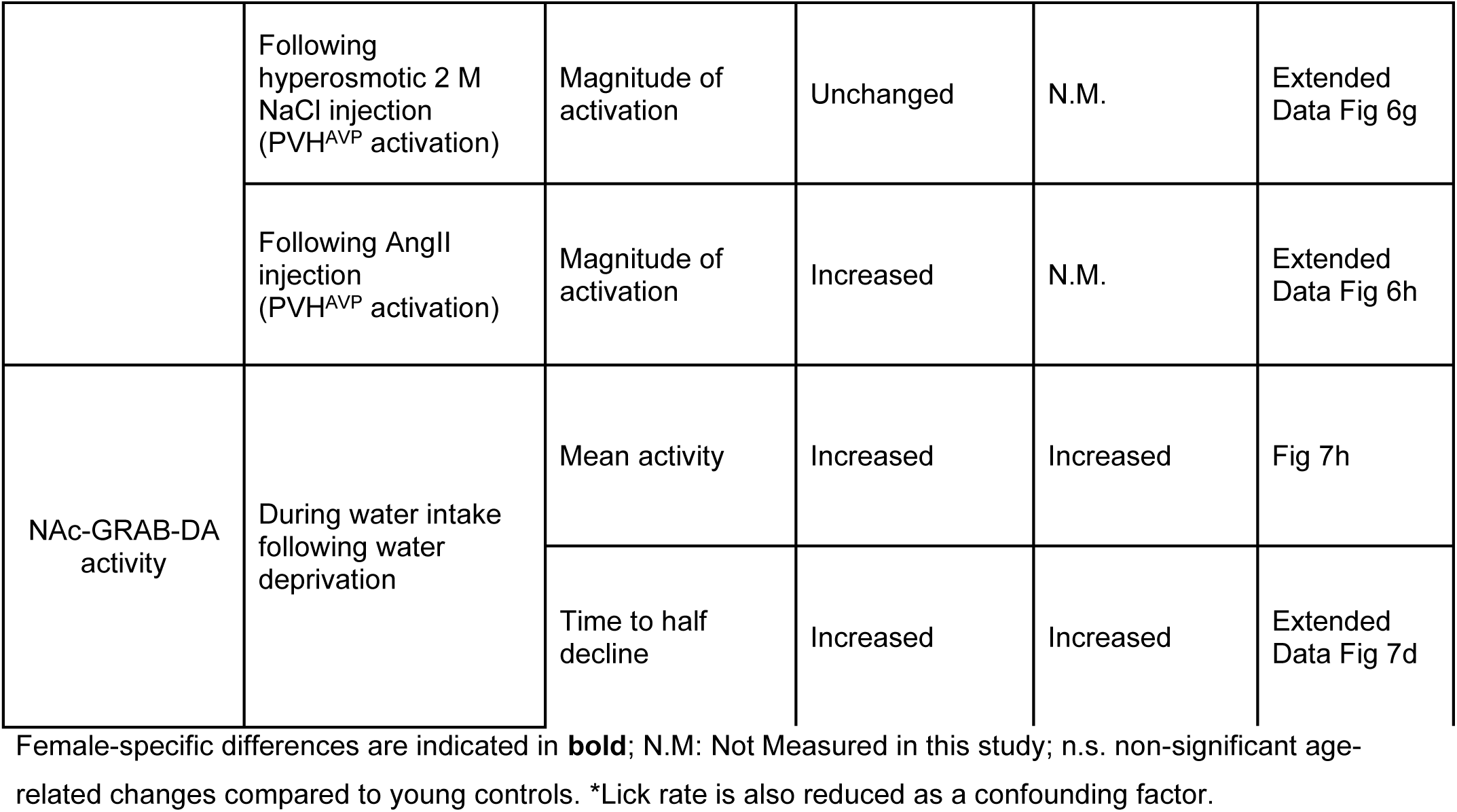
Age-related changes in male and female mice from our study.

### Aged mice show unexpectedly enhanced thirst after water deprivation

A surprising result from our experiments was that aged male and female mice consumed more water than young animals after 24-hour water deprivation (**Fig 5d**). This consistent result was accompanied by increased motivation to work for water (**Fig 7b-d**) and increased dopamine release in the NAc in response to water consumption (**Fig 7h**), suggesting that aged mice have greater activation of the thirst circuit after 24-hour deprivation. This is at odds with the general observation that aged animals show reduced thirst, and indeed our finding that aged mice are chronically dehydrated (**Fig 1c**). It also appears inconsistent with our finding that aged animals do not overdrink in response to isolated blood hyperosmolality or hypervolemia (**Fig 8b, i; Extended Data Fig 8g, k**).

How can we explain these findings? The simplest explanation is that 24-hour water deprivation causes substantially greater dehydration in aged mice than young animals. This would be predicted from the fact that aged animals have decreased AVP function and impaired urine concentrating ability (**Fig 1d-g**), and, for this reason, it was surprising that we observed similar blood osmolality in young and old animals after 24-hour water deprivation (**Fig 5a**). We also found that the reduction of blood volume by 24-hour water deprivation is similar in young and aged animals (**Extended Data Fig 8c**). However, if thirst is influenced by the duration of blood hyperosmolality, then the fact that aged animals likely become dehydrated faster in response to water deprivation may be important. Addressing these questions will require measuring the time course of blood osmolality and volume following water deprivation in young and old animals and correlating this with behavioral and neural endpoints.

### Limitations of this study

Although our study provides a comprehensive analysis of how the fluid homeostasis system changes with aging, there are technical and conceptual limitations that we want to emphasize. First, our experiments rely extensively on fiber photometry, which can measure relative changes in calcium dynamics on a short timescale (tens of minutes) but is poorly suited to measuring changes in absolute activity levels over days or longer. Second, while we observe several interesting, age-dependent differences in the temporal coordination between eating and drinking, both at the level of behavior and neural dynamics, we cannot draw conclusions about how these changes in microstructure relate to longer term changes in fluid homeostasis. Third, there are ceiling effects in some behavioral assays (e.g. the rate of water ingestion following optogenetic stimulation of SFO-Glut neurons). Thus, differences water intake could reflect physical constraints rather than differences in thirst alone.

## METHODS

### Mouse strains

The C57BL/6J (Jackson laboratory, 000664) mouse strain was used. Mice were housed or aged (for aged mice) in the temperature- and humidity-controlled facilities with 12-hr light-dark cycle and *ad libitum* access to water and standard chow (PicoLab Rodent Diet 20 #5053). Avp-IRES2-Cre (JAX #023530) mice were obtained from the Jackson laboratory and bred with wild-type mice to generate AVP-Cre/+ mice for experimental use. Aged C57BL/6JN mice from the NIA aged rodent colony were used for Extended Data Fig 1a, 5a-b and 8a-c.

### Plasma and urine osmolality

For plasma, blood was collected from cheek vein into the lithium-heparin-coated tube (RAM scientific) and 1,000xg for 10 minutes, and the supernatant was further centrifuged at 10,000 xg for 10 minutes and to remove platelets. Urine was collected by spontaneous voiding or gentle massage of the abdomen, and centrifuged at 1,000 xg for 5 minutes and the supernatant was diluted by 5 times in water. Osmolality was measured using the Vapro 5600 vapor-pressure osmometer (Wescor Instruments).

### dDAVP (Desmopressin) injection

dDAVP (Desmopressin, Cayman Chemical #17348) was dissolved in DMSO to make a 3.35 mg/mL stock, kept frozen -20’C and diluted in PBS on the same day for subcutaneous injection at 300 uL s.c./20g BW (1 mg s.c./kg BW). Urine was collected in the baseline state, 2 hours and 4 hours after dDAVP injection for osmolality measurements.

### Urine AVP and Plasma copeptin measurements

Urine AVP levels were quantified using an Arg8-Vasopressin ELISA kit (Enzo, ADI-900-017A), and plasma copeptin levels were measured using an ELISA kit (Cloud-Clone Corp., CEA365Mu). For both assays, optical density at 450 nm was measured using a Biotek H4 Plate Reader.

### Blood volume

Relative changes in blood volume within the same cohort were estimated by measuring plasma protein concentration using a bicinchoninic acid (BCA) protein assay (Pierce, 23225). Because plasma protein concentration is inversely proportional to blood volume, changes in protein concentration before and after stimulation were used as an index of relative blood volume changes.

### Void spot assay and measurement of urine volume

Urine volume was measured by spontaneous voiding on a clean paper (Blick Cosmos blotting paper, 10422-1005). A clean housing cage was removed of all bedding materials, food and water, and prepared with a clean paper (16.0 cm x 30.1 cm) cut and taped on the bottom to cover the entire floor, and 2 pieces of bedding and 1 piece of chow on the paper. An individual mouse was placed in the cage and left in the dark, sound-attenuating PVC cabinet (Med Associates Inc, ENV-018V) for 4 hours from 10 AM to 2 PM. After 4 hours, the mouse was returned to its home cage and the paper was imaged by a gel imaging system (ChemiDoc^TM^ MP, Universal Hood II, BioRad) with the Krypton detection setting. The imaged void area was analyzed using the Void Whizzard plugin^1^ on ImageJ. The size of the void area on the paper was converted to urine volume by a standard calibration curve with a series of known volumes of urine which had a linear relationship with the size of the urine spot. A spot size corresponding to 0.5 uL of urine or smaller was excluded to avoid nonspecific marks.

### *ad libitum* feeding and drinking behavior

Baseline behavior and behavior only cohort with a thirst stimulus was measured using the Coulbourn Habitest system® (Harvard Apparatus). For food consumption behavior, regular pellets (BioServ Dustless Precision Pellets, F0071) were dispensed from an automatic feeder operated by an optical sensor. The weight of individual pellet was measured by weighing 50 pellets 5 times independently, and was 18.0 mg (mean) +/- 0.3 mg (standard deviation). For water consumption behavior, contact lickometers were manually built to detect and record individual licks^2^. To measure the volume of individual water licks, the water bottle was weighed at the beginning and the end of an overnight session of *ad libitum* feeding and drinking in the behavioral chamber, and the volume of consumed water was divided by the total number of licks, yielding 1.21 μL (mean) +/- 0.02 μL (standard deviation). For baseline behavior, mice were given *ad libitum* access to pellets and water to measure baseline food and water intake behavior for 24 hours. For prandial thirst behavior, mice were fasted overnight with access to water and given *ad libitum* access to pellets for 30 minutes with the water spout blocked. After 30 minutes, the pellet dispenser was stopped and the water spout was exposed for water lick measurement for the following 30 minutes. For baseline behavior and prandial thirst behavior, crumbs generated during pellet consumption were collected on a tray underneath the behavioral chamber and were weighed after the experiment for subsequent correction of total food intake. For water deprivation thirst behavior, mice were water deprived for 24 hours in their home cage with access to chow prior to the water lick measurement in the behavior chamber. For osmotic thirst behavior, mice were intraperitoneally injected with 2 M NaCl. For hypovolemic thirst behavior, mice were subcutaneously injected with 40% of polyethylene glycol in their shoulder. All mice were habituated to the behavioral chamber, lickometer and/or feeder one overnight prior to the experiments. All behavioral recordings started after mice were initially acclimated to the behavioral chamber for 15 minutes. Mice had at least 24 hours of recovery after each experiment before initiation of a different thirst paradigm.

### Behavioral analysis

Behavioral data was collected by Graphic State software and subsequently analyzed using custom-written python scripts. A meal was defined as 50 mg of food consumption spaced by 5 minutes or less^3,4^. Water lick bouts were defined as 10 or more water licks spaced by 1 second or less^5^.

### Lever press operant task

A lever-press operant task was conducted using a Coulbourn Habitest system® (Harvard Apparatus). A water spout was connected to a solenoid valve (LEE company #LHLA2431111H) via a custom-designed circuit. Pressing the designated active lever (but not the inactive lever) triggered the valve to open for 50 milliseconds, delivering a water reward of about 6.49 μL. Three reinforcement schedules were employed: In Fixed Ratio 1 (FR1), a single press on the active lever resulted in one valve opening event and water delivery. In Fixed Ratio 5 (FR5), five consecutive presses on the active lever were required for a single valve opening event and water delivery. In Progressive Ratio (PR), the number of active lever presses needed for a valve opening event and water delivery progressively increased in a fixed-increment sequence (in multiples of 3: 3, 6, 9, 12, …).

Mice were initially trained for the task using the FR1 schedule for five consecutive nights before FR1 experiments. Following FR1 experiments, mice underwent training on the FR5 schedule for three nights prior to FR5 experiments. Finally, the PR schedule was implemented. Overnight training was performed with *ad libitum* access to chow. For all lever press experiments, mice were water deprived for 24 hours prior to the experiment. Following water deprivation, mice were acclimated to the behavioral chamber for 15 minutes. During acclimation, the wall containing the levers and water spout remained occluded to prevent interaction with the operant conditioning apparatus After 15 minutes, experiments started with the levers and water spout exposed. Mean number of lever press was plotted by aligning to the first lever press.

### Stereotaxic surgery

For PVH-AVP fiber photometry recordings, AVP-Cre/+ mice were injected with 150 nL of AAV1.CAG.flex.GCaMP6s (Janelia Vector Core, 4.73×10^13^ vg/mL) into the PVH (−0.7 mm AP, +0.3 mm ML, −4.7 mm DV). In the same surgery, an optical fiber (Doric Lenses, 0.4 mm inner diameter and 5.4−6.0 mm length, MFC_400/430-0.48) was implanted in the same coordinate as the injection site. For SFO-Glut fiber photometry recordings, wild-type mice were injected with 200 nL of AAV9.CaMKII.GCaMP6s.WPRE (Janelia Vector Core, 2.17×10^14^ vg/mL) freshly diluted 1/10 with sterile PBS into the SFO (-0.65 mm AP, 0 mm ML, −2.7 mm DV). In the same surgery, an optical fiber (Doric Lenses, 0.4 mm inner diameter and 3.0−3.5 mm length, MFC_400/430-0.48) was implanted at -0.65 mm AP, 0 mm ML, −2.85 mm DV in young animals and −0.65 mm AP, 0 mm ML, −2.9 mm DV in aged animals.

For NAc-DA fiber photometry recordings, wild-type mice were injected with 200 nL of AAV9.hSyn.GRAB-DA3m (WZ Biosciences, 5.80×10^13^ vg/mL) into the NAc shell (+1.0 mm AP, +1.4 mm ML, −4.5 mm DV). In the same surgery, an optical fiber (Doric Lenses, 0.4 mm inner diameter and 5.0−5.4 mm length, MFC_400/430-0.48) was implanted in the same coordinate as the injection site. For SFO optogenetics experiments, wild-type mice were injected with 150 nL of AAV5.CaMKIIa.hChR2 (Janelia Vector Core, 7.36×10^12^ vg/mL) into the SFO (−0.65 mm AP, 0 mm ML, −2.65 mm DV). In the same surgery, an optical fiber (Thorlabs, 200 µm) was manually trimmed and polished to 3.0−3.5 mm length and implanted 0.25 mm above the injection site.

Following surgical implantation, all mice were allowed a recovery period of two weeks and were habituated in the behavioral chamber overnight with a lickometer prior to fiber photometry or optogenetic experiments.

### Fiber photometry recording and analysis

#### Tethering and light delivery

Implanted mice were secured with a patch cable (Doric Lenses, MFP_400/460/900-0.48_2m_FCM-MF2.5). Continuous excitation light was provided by two light-emitting diodes (LEDs): a 6 mW blue LED with a wavelength of 470 nm and a UV LED with a wavelength of 405 nm. A multichannel hub from Thorlabs served as the power source for the LEDs. To differentiate between signals, these LEDs were modulated at distinct frequencies of 211 Hz and 511 Hz, respectively. A filtered minicube (Doric Lenses, FMC6_AE(400-410)_E1(450-490)_F1(500-540)_E2(550-580)_F2(600-680)_S) ensured that only specific wavelengths reached the implant. The filtered light was then channeled via optical fibers (Doric Lenses, MFC_400/430-0.48) to the implanted device.

#### Signal acquisition and processing

Light signals originating from either GCaMP or GRAB-DA, along with UV isosbestic point signals, were simultaneously collected through the same optical fibers using a femto-watt silicon photoreceiver (Newport, 2151). The collected digital signals were amplified using a lock-in amplifier to enhance the desired signal and eliminate background noise. Then, they were demodulated using their corresponding modulation frequencies (211 Hz and 511 Hz) to separate the signals originating from the two LEDs. Finally, an RZ5P processor (Tucker-Davis Technologies) processed the signals for further analysis. The processed data from the fiber photometry recordings were collected and stored using Synapse software (TDT).

#### Lickometer recording and data analysis

A custom-designed lickometer based on the work of Slotnick (2009)^2^ was employed to detect individual water lick events. These lick events were time-stamped and recorded by the TDT software throughout the fiber photometry recording session. After data exportation via Browser (TDT), custom Python scripts were used to analyze the data following downsampling at a rate of 1/125.

#### Data processing

GCaMP or GRAB-DA signals were normalized by UV signals, and z score was used for plots and subsequent analysis (z = (x-μ)/σ, z is the raw signal, μ is the population mean, and σ is the population standard deviation). z score values were smoothened by a factor of 101 using the Hanning window for calculating half time for GCaMP activation and linear regression fitting during SFO-Glut GCaMP inhibition. For activation during chow consumption or following 2 M NaCl injection, the time required for the GCaMP signal to reach 50% of its peak value relative to baseline (T_GCaMP half activation_). For activation following PEG injection, which produced slower and more gradual responses, the time required for the GCaMP signal to reach 33% of peak value relative to baseline was used (T_GCaMP 1/3 activation_). For water drinking following chow intake or water deprivation, the half-time for the GCaMP signal to reach the transition point from the initial rapid SFO-Glut inhibitory phase to the subsequent steady phase, determined by linear regression fitting, was quantified. For water drinking following 2 M NaCl or PEG injection, the time for GCaMP inhibition to decline from its peak to 50% (T_GCaMP half inhibition_) or 67% (T_GCaMP 1/3 inhibition_) of peak relative to baseline was used, respectively.

#### Recording of feeding behavior

For prandial thirst, animals were fasted overnight and given access to standard chow (PicoLab Rodent Diet 20, 5053) without water for the first 30 minutes, followed by chow removal and water access for the next 30 minutes. Chow was weighed before and after the experiment and fallen chow crumbs were weighed to obtain true chow consumption. Chow intake was video recorded with an infra-red camera and iSpy software. Individual chow bite events were scored using the BORIS software^6^. To note, standard chows were provided on the floor of the chamber instead of utilizing the automated pellet dispenser. This modification was necessary due to physical incompatibility between the fiber optic implants on the mouse head and the design of the feeding trough associated with the automated pellet dispenser.

#### Reagent preparation

2 M NaCl was prepared in water and intraperitoneally injected at 250 μL/35 g body weight. 40% polyethylene glycol (PEG 400, Sigma #807485) was prepared in water and subcutaneously injected in the shoulder at 300 μL/20 g body weight. Angiotensin II (AngII, Anaspec, AS-20633) stock was prepared in PBS (10 μg/μL), kept in -80°C in aliquots, and prepared fresh on the day of experiment for subcutaneous injection at 3.0 mg/kg body weight in PBS.

### Optogenetic stimulation

For optogenetic stimulation of SFO-Glut neurons, mice were either well-hydrated or water deprived for 24 hours in their home cages, acclimated to the behavioral chamber for 15 minutes, then provided access to water for 15 min with or without optical stimulation (15 mW, 20 Hz) using a DPSS 532 nm laser (Shanghai Laser and Optics Century). Water lick events were recorded for 30 minutes. For terminal SFO-Glut stimulation for c-fos immunohistochemistry, mice were photo-stimulated for 15 minutes (15 mW, 20 Hz), waited for 90-120 minutes, then transcardiac perfused with saline followed by 4% paraformaldehyde for collection of the brain.

### Immunohistochemistry

At the end of their experiments, mice were transcardiac perfused with saline followed by 4% paraformaldehyde (PFA). Brain was collected and incubated in 4% PFA overnight at 4°C, followed by incubation in 30% sucrose in PBS at 4°C for 1-3 days, freezing in the OCT cryoprotectant compound (Tissue-Tek) and storage at -80°C. Frozen brains were sectioned in 30-40 μm thickness and incubated in normal goat serum (NGS) for 1 hour in RT, stained with primary antibodies for one overnight at 4°C, washed in 10% Triton-X in PBS three times, stained with secondary antibodies for 1 hour in RT, and washed in PBS three times before mounting onto the glass slide for microscopy. Brain sections were imaged using Zeiss LSM 700 confocal microscope and Nikon Eclipse Ti2 microscope. For GCaMP and GRAB-DA, chicken anti-GFP polyclonal antibody (Abcam, ab13970, 1:2000) and goat anti-chicken IgY (H+L) antibody, Alexa Fluor™ 488 (Thermo Fisher, A-11039, 1:1000) was used primary and secondary staining, respectively. For AVP, rabbit anti-AVP polyclonal antibody (Sigma-Aldrich, AB1565, 1:1000) and goat anti-rabbit IgG (H&L) antibody, Alexa Fluor® 568 (abcam, ab175471, 1:1000) was used for primary and secondary staining, respectively.

### Statistics and figures

Statistical analyses were performed using the Prism software. Schematics for experimental procedures were generated using Biorender.

## Author contributions

H.J., Z.A.K. and J.L.G. conceived the project and designed experiments. H.J. led and performed experiments. H.J, Z.A.K. and J.L.G. analysed and interpreted data. H.J., A.B.S., U.D. and J.H.W. collected plasma and urine samples and measured the parameters. H.J., A.B.S. and U.D. performed behavioral experiments. H.J., A.B.S. and J.H.W. performed the photometry experiments. H.J. and A.B.S. performed the optogenetic experiments. H.J. performed all the photometry and optogenetic surgeries. H.J., A.B.S. and J.H.W. performed histology. H.J, Z.A.K. and J.L.G wrote the manuscript.

### Acknowledgements

We thank members of the Garrison and Knight labs for constructive discussions; Jonathan Riggle for providing the video-recording setup for pellet feeding; Sarah Shehata for assistance with analyzing chow feeding videos; Courtney Hudson-Paz and Jamie S. Ahn for assistance with mouse tissue collection and histology; Cyrus Ghiassi, Alba Villagrana and Elizabeth Poznyakov for assistance with imaging void spot papers. This work was funded in part by the NIH (R35 GM119828 and R35 GM145305 to J.L.G., T32AG000266 and F32AG063488 to H.J.), and the Glenn Foundation for Medical Research (to H.J.). Zachary A. Knight is a Howard Hughes Investigator.

