## Supplemental Figures for "Dysregulation of the fluid homeostasis system by aging"

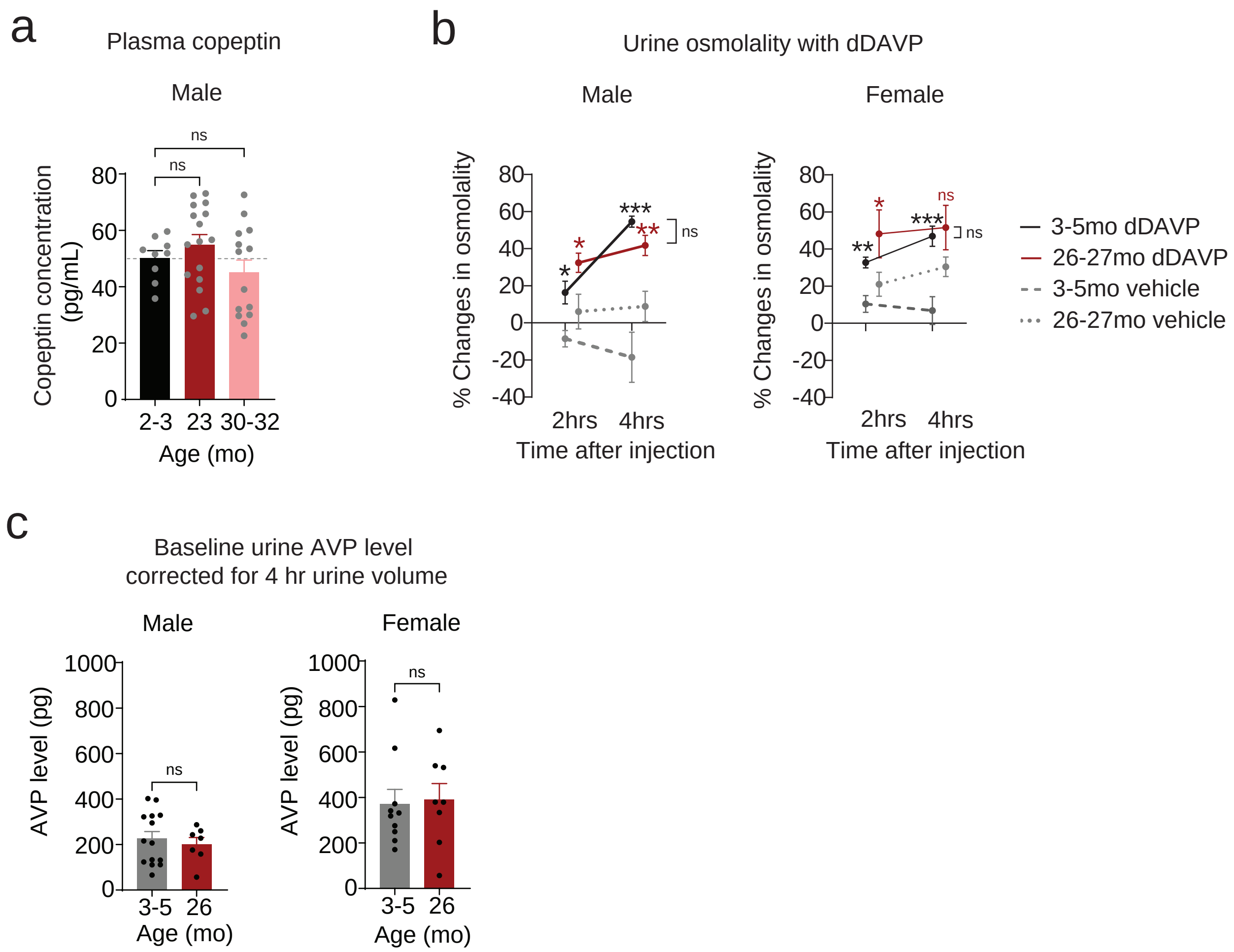


**Extended Data Fig 1. Aged mice have renal dysfunction yet unaltered circulating copeptin levels**
**a.** Circulating copeptin in plasma in 2-3, 23, 30-31 month-old male mice. n.s., non-significant by one-way ANOVA, P=0.005 by Brown-Forsythe test. n.s., non-significant by Dunnet's multiple comparisons test between age groups.
**b.** Relative changes in urine osmolality from the baseline value, 2 hrs and 4 hrs after dDAVP (1 mg/kg BW) injection in 3-5 and 26-27 month-old male (left) and female (right) mice. n.s., non-significant, *P < 0.05, **P<0.01 and ***P<0.001 by Fisher's LSD test, compared to the vehicle control 2 hrs and 4 hrs after injection, respectively, or between designated pairs. For males, ***P<0.001 for main effects of dDAVP [F(1,26)=37.29], non-significant for main effects of age [F(1, 26)=0.7052] and *P<0.05 for their interactions [F(1,26)=5.348] by 2-way ANOVA. For females, ***P<0.001 for main effects of dDAVP [F(1,24)=16.82], and non-significant for main effects of age [F(1, 24)=3.611] and for their interactions  [F(1,24=1.613] by 2-way ANOVA.
**c.** Baseline urine AVP level normalized to urine volume collected for 4 using the void spot assay. n.s., non-significant by Welch's t-test.

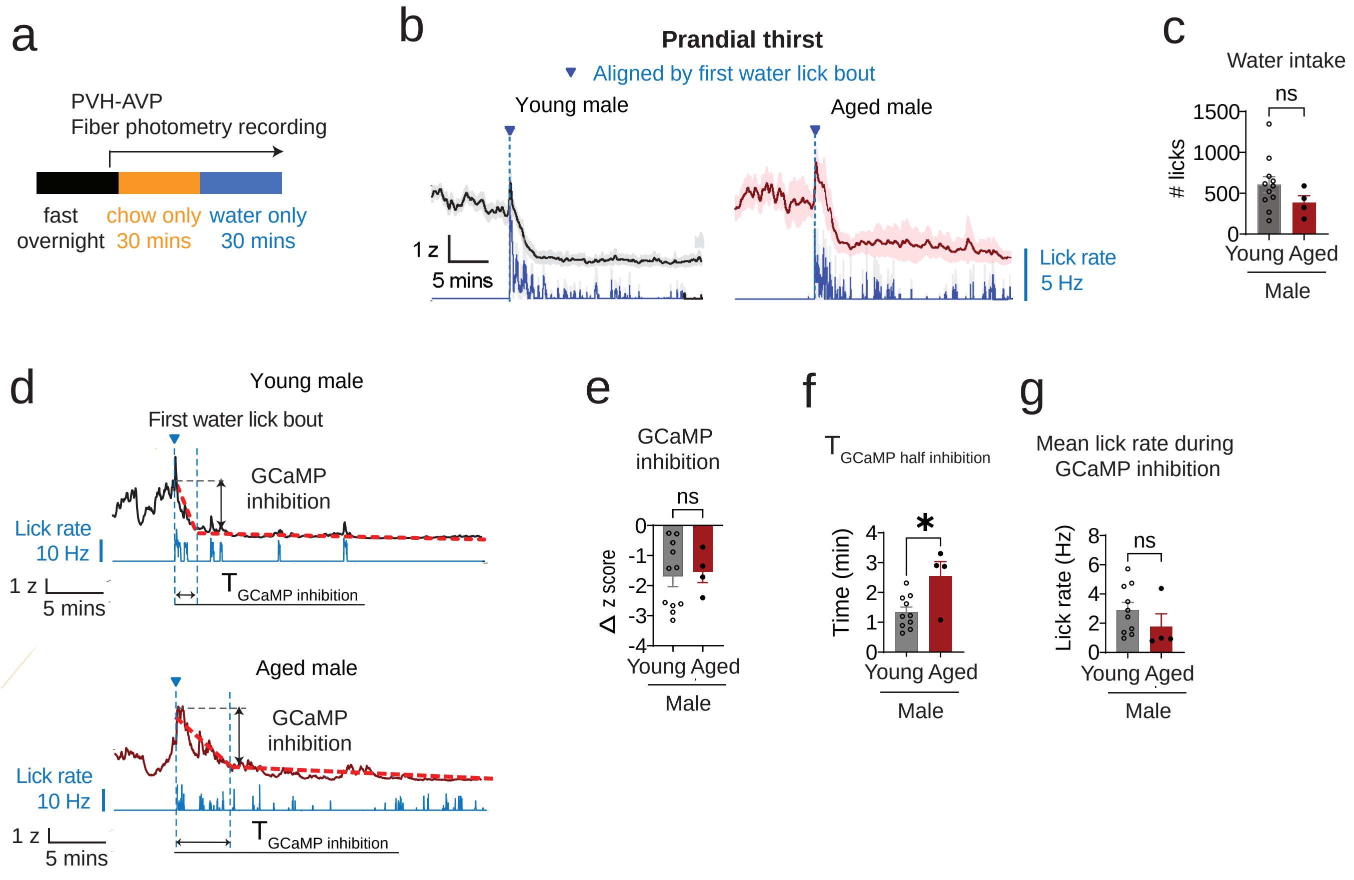


**Extended Data Fig 2. PVH$^{AVP}$ recording during drinking after eating**

**a.** Experimental paradigm for PVH$^{AVP}$ fiber photometry recording during 30 minute chow intake following overnight fasting, followed by 30 minutes of water intake.

**b.** Mean lick rate (blue) ± s.e.m. and mean z-score trace of GCaMP fluorescence ± s.e.m. in young (3-4 mo, black) and aged (24-26 mo, magenta) male mice, aligned to the first water lick bout (blue dotted line).

**c.** Total number of water licks during 25 minutes following 30 minutes of chow intake. n.s., non-significant by unpaired t-test.

**d.** Representative lick rate (blue) and GCaMP fluorescence (z-score) traces from a 4-month-old (top; black) and a 26-month-old (bottom; magenta) male mouse, aligned to the first water-lick bout. Red dotted lines represent linear regression fits for the fast PVH$^{AVP}$ inhibition phase and the subsequent steady phase.

**e.** Magnitude of PVH$^{AVP}$ inhibition during the fast inhibition phase as indicated in **d.** n.s., non-significant by unpaired t-test.

**f.** Half time for GCaMP signals to reach the transition from the fast PVH$^{AVP}$ inhibition phase to the steady phase as indicated in **d.** *P<0.05 by unpaired t-test.

**g.** Mean lick rate during the fast PVH$^{AVP}$ inhibition phase. n.s., non-significant by unpaired t-test.

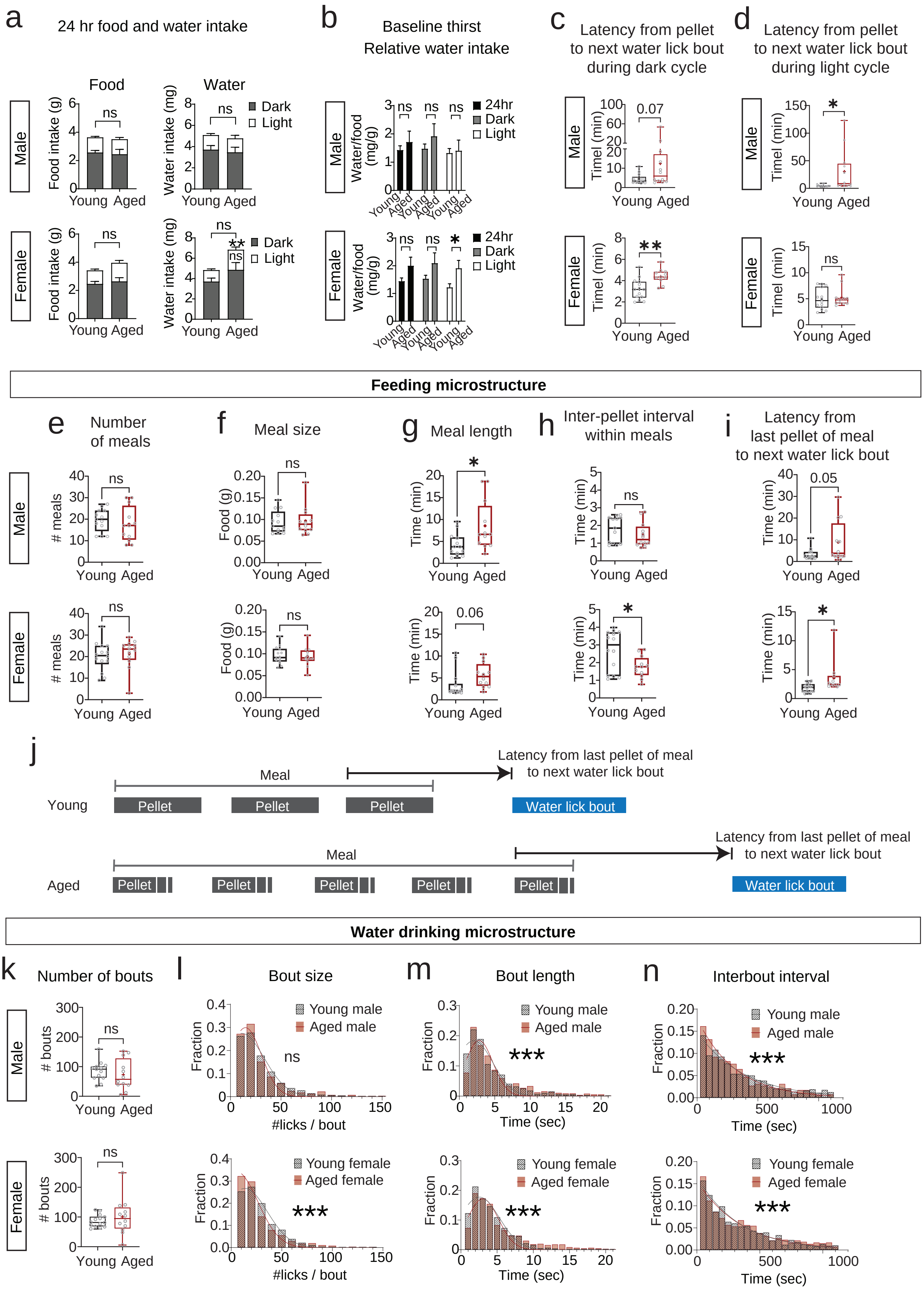

a
24 hr food and water intake
Food
Water
Male
Female
Food intake (g)
Water intake (mg)
Dark
Light
ns
Young Aged
b
Baseline thirst
Relative water intake
Water/food (mg/g)
24hr
Dark
Light
c
Latency from pellet to next water lick bout during dark cycle
0.07
d
Latency from pellet to next water lick bout during light cycle
Timel (min)
Feeding microstructure
e
Number of meals
### meals
f
Meal size
Food (g)
g
Meal length
Time (min)
0.06
h
Inter-pellet interval within meals
i
Latency from last pellet of meal to next water lick bout
0.05
j
Meal
Latency from last pellet of meal to next water lick bout
Young
Aged
Pellet
Water lick bout
Water drinking microstructure
k
Number of bouts
### bouts
l
Bout size
Fraction
#licks / bout
Young male
Aged male
Young female
Aged female
m
Bout length
Time (sec)
n
Interbout interval

**Extended Data Fig 3. Baseline food and water intake**

**a.** Baseline food and water intake for 24 hours. (Left) total food intake during 12 hours of dark cycle (gray) and 12 hours of light cycle (white) in young (3-5 mo) and aged (26-27 mo) male and female mice. (Right) total water intake during 12 hours of dark cycle (gray) and 12 hours of light cycle (white) in male and female mice.

**b.** Water intake relative to food intake during 24 hours (black), 12 hours of dark cycle (gray) and 12 hours of light cycle (white) in male and female mice. n.s., non-significant, *P<0.05, and **P<0.01 by Welch's t-test.

**c.** Latency from each pellet retrieval to the first following water lick bout.E ach dot represents the median value during the dark cycle for an individual mouse within each sex and age group. **P<0.01 by unpaired t-test.

**d.** Latency from each pellet retrieval to the first following water lick bout. Each dot represents the median value during the light cycle for an individual mouse within each sex and age group. *P<0.05 by unpaired t-test.

**e.** Number of meals during the dark cycle. n.s., non-significant,by unpaired t-test.

**f.** Amount of food intake per meal during the dark cycle. Each dot represents the median value for an individual mouse within each sex and age group. n.s., non-significant by unpaired t-test.

**g.** Duration of meal during the dark cycle. Each dot represents the median value for an individual mouse within each sex and age group.*P<0.05 and **P<0.01 by Welch's t-test.

**h.** Time interval between pellet retrieval within a meal during the dark cycle. Each dot represents the median value for an individual mouse within each sex and age group. *P<0.05 by Welch's t-test.

**i.** Latency from the last pallet of a meal to the first following water lick bout during the dark cycle. Each dot represents the median value for an individual mouse within each sex and age group. *P<0.05 by Welch's t-test.

**j.** Diagram describing differences in the feeding microstructure between young and aged mice. Broken boxes for pellet in aged animals represent pellet spills and drops.

**k.** Number of water lick bouts during the dark cycle. n.s., non-significant by Welch's t-test.

**l** Histogram of the size of water lick bouts pooled across sex and age groups during the dark cycle. n.s., non-significant and ***P<0.001 by Kolmogorov-Smirnov test.

**m.** Histogram of the length of water lick bouts pooled across sex and age groups during the dark cycle ***P<0.001 by Kolmogorov-Smirnov test.

**n.** Histogram of the interbout interval of water lick bouts pooled across sex and age groups during the dark cycle. ***P<0.001 by Kolmogorov-Smirnov test.

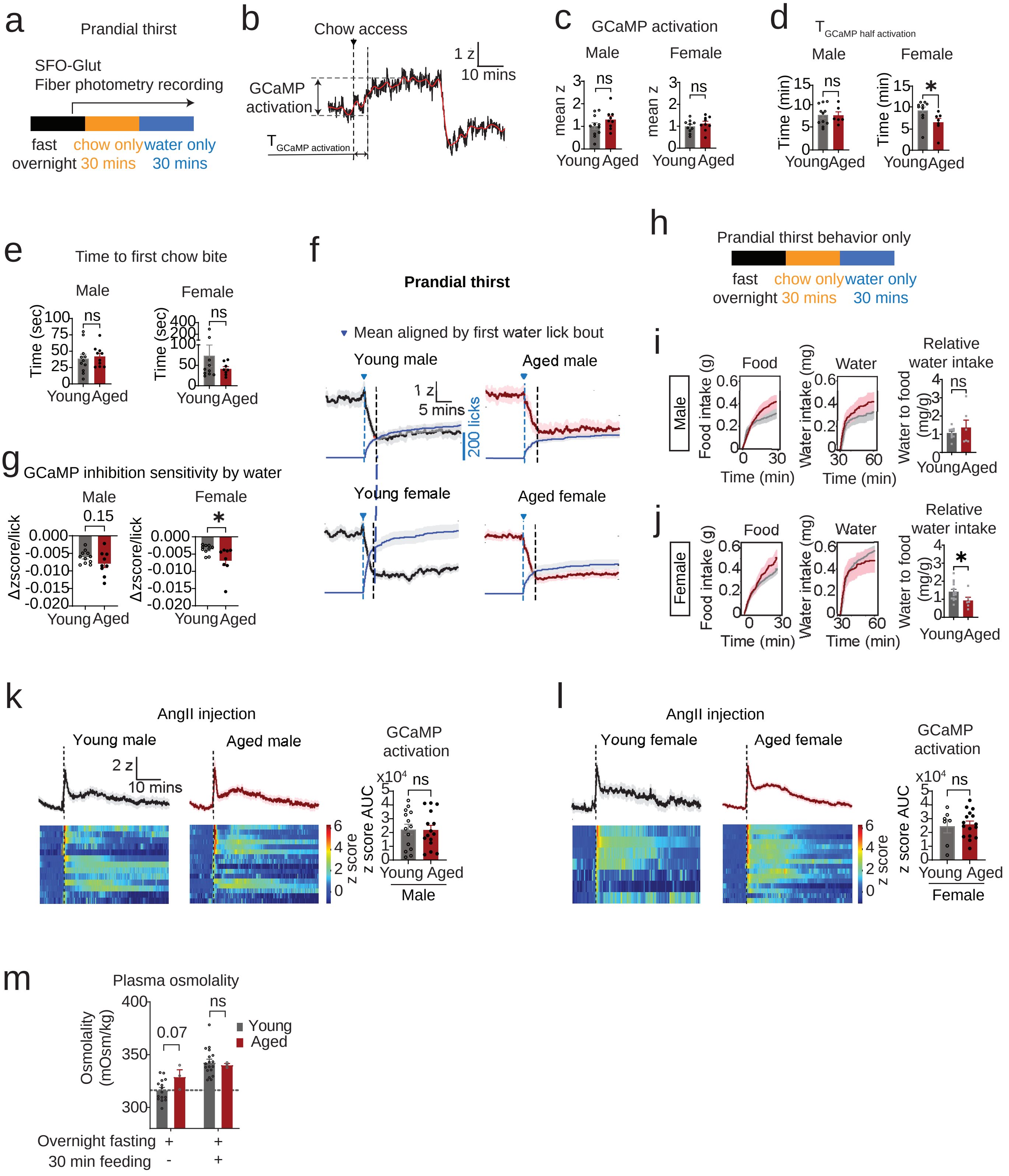
a
Prandial thirst
SFO-Glut
Fiber photometry recording
fast overnight
chow only 30 mins
water only 30 mins
b
Chow access
1 z
10 mins
GCaMP activation
T GCaMP activation
c
GCaMP activation
Male
Female
mean z
ns
Young Aged
d
T GCaMP half activation
Male
Female
Time (min)
ns
*
Young Aged
e
Time to first chow bite
Male
Female
Time (sec)
ns
Young Aged
f
Prandial thirst
Mean aligned by first water lick bout
Young male
Aged male
1 z
5 mins
200 licks
Young female
Aged female
g
GCaMP inhibition sensitivity by water
Male
Female
0.15
*
Δzscore/lick
Young Aged
h
Prandial thirst behavior only
fast overnight
chow only 30 mins
water only 30 mins
i
Male
Food
Water
Relative water intake
Food intake (g)
Water intake (mg)
Water to food (mg/g)
Time (min)
ns
Young Aged
j
Female
Food
Water
Relative water intake
Food intake (g)
Water intake (mg)
Water to food (mg/g)
Time (min)
*
Young Aged
k
AngII injection
Young male
Aged male
2 z
10 mins
z score
GCaMP activation
z score AUC
x10^4
ns
Young Aged
Male
l
AngII injection
Young female
Aged female
z score
GCaMP activation
z score AUC
x10^4
ns
Young Aged
Female
m
Plasma osmolality
Osmolality (mOsm/kg)
0.07
ns
Young
Aged
Overnight fasting + +
30 min feeding - +

**Extended Data Fig 4. $SFO^{Glut}$ recording during drinking after eating and AngII injection**
**a.** Behavioral paradigm for $SFO^{Glut}$ fiber photometry recording during 30 minute chow intake after overnight fasting followed by 30 minutes of water intake.
**b.** Representative z-score trace of GCaMP fluorescence during 30 minutes of chow intake followed by 30 minutes of water drinking in a 3-month-old male mouse after overnight fasting. The red line indicates a smoothened trace; black lines denote chow access, the magnitude of GCaMP activation and the time to half-maximal activation
**c.** Magnitude of GCaMP activation, as shown in **b**. ns., non-significant.
**d.** Time for GCaMP signals to rise from baseline to the half peak z-score of GCaMP fluorescence, as shown in **b**. *P<0.05 and ns., non-significant by unpaired t-test.
**e.** Latency from the chow access to the first chow bite. ns., non-significant.
**f.** Mean cumulative water licks (gray) and mean GCaMP fluorescence (z-score) in young (black) and aged (magenta) male and female mice, aligned to the first water-lick bout (blue dotted line with triangle).
**g.** Magnitude of GCaMP inhibition per water lick during the fast inhibition phase. *P<0.05 and ns., non-sgnificant by unpaired t-test.
**h.** Behavioral paradigm for 30 minutes of chow intake after overnight fasting followed by 30 minutes of water intake.
**i.** (Left) Cumulative pellet intake after overnight fasting, (middle) cumulative number of water licks after 30 minutes of refeeding, and (right) water intake normalized to food intake in young (2-5 mo) and aged (26-27 mo) male mice from the behavior only cohort. ns., non-significant by unpaired t-test.
**j.** (Left) Cumulative pellet intake after overnight fasting, (middle) cumulative number of water licks after 30 minutes of refeeding, and (right) water intake normalized to food intake in young (2-5 mo) and aged (26-27 mo) female mice from the behavior only cohort. *P<0.05 by unpaired t-test.
**k.** (Top) Mean z-score trace of GCaMP fluorescence in young (3-5 mo; black) and aged (23-27 mo; magenta) male mice following subcutaneous angiotensin II (AngII) injection (black dotted line). (Bottom) Heatmaps of individual z-score traces. (Right) Area under the curve of GCaMP fluorescence during the 45 minutes following AngII injection. ns, non-significant by unpaired t-test.
**l.** (Top) Mean z-score trace of GCaMP fluorescence in young (2-5 mo, black) and aged (24-30 mo, magenta) female mice following subcutaneous angiotensin II (AngII) injection (black dotted line). (Bottom) Heatmaps of individual z-score traces. (Right) Area under the curve of GCaMP fluorescence during the 45 minutes following saline injection. ns, non-significant by unpaired t-test.
**m.** Plasma osmolality measured before and after 30 minutes of chow intake in overnight-fasted young (2-6 mo) and aged (28-33 mo) male mice.

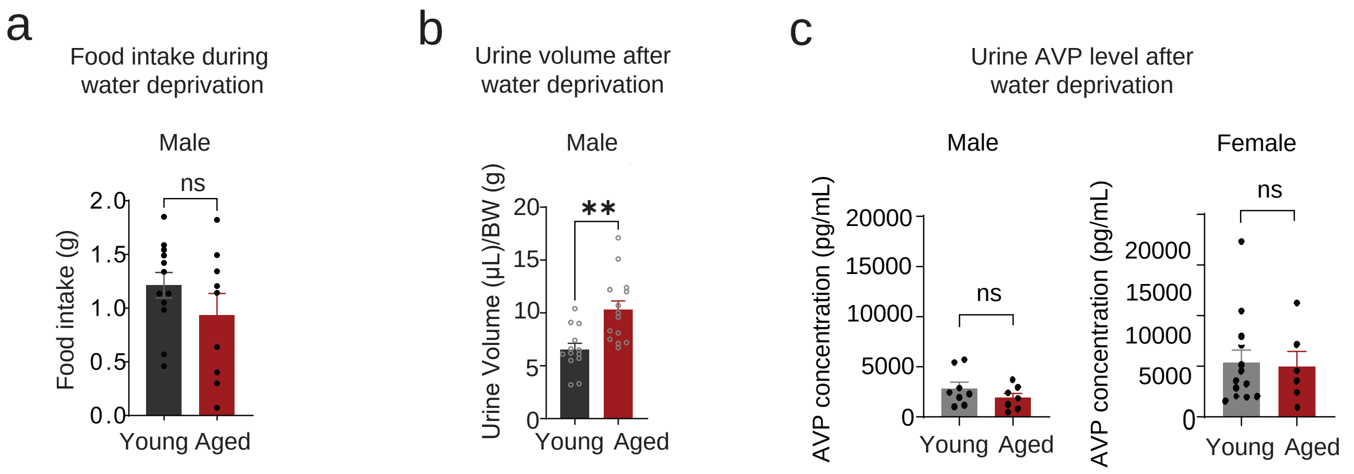


**Extended Data Fig 5. Age-related effect of water deprivation on food intake and urine volume.**
**a.** Food intake during water deprivation in the dark cycle in young (3-5 mo) and aged (25 mo) male mice. n.s., non-significant by Welch's t-test..
**b.** Urine volume for 4 hours following 24 hours of water deprivation in young (3-5 mo) and aged (25 mo) male mice. **P<0.01 by Welch's t-test
**c.** Urine AVP levels in young (3-5 mo) and aged (26-27 mo) male and female mice following 24 hours of water deprivation. ns, non-significant by Welch's t-test.

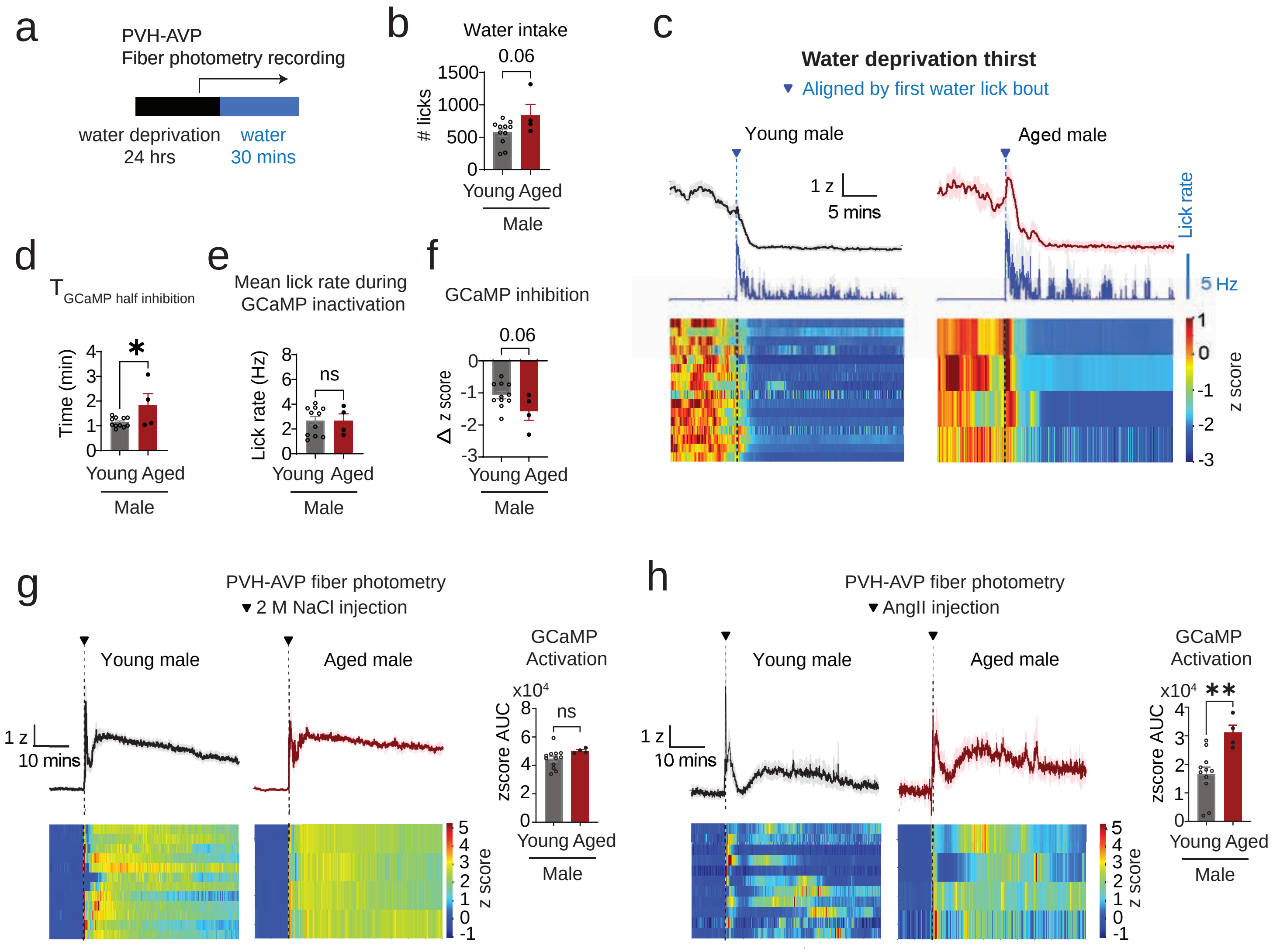


**Extended Data Fig 6. PVH$^{AVP}$ recording during drinking after water deprivation and during acute injection of hypertonic saline and AngII.**

**a.** Experimental paradigm of PVH$^{AVP}$ recording during drinking after water deprivation.

**b.** Total number of water licks during 25 minutes in the young (3-4 mo) and aged (24-26 mo) PVH$^{AVP}$ fiber photometry mice.

**c.** (Top) Mean lick rate (blue) and mean z-score trace of GCaMP fluorescence in young (black) and aged (magenta) male and female mice, aligned to the first water lick bout (blue dotted line). (Bottom) Heatmaps of individual z-score traces in each group. Black dotted lines represent the first water lick bout.

**d.** Half time for GCaMP signals to reach the transition from the fast PVH$^{AVP}$ inhibition phase to the steady phase. *P<0.05 by unpaired t-test.

**e.** Mean lick rate during the fast PVH$^{AVP}$ inhibition phase. ns, non-significant by unpaired t-test.

**f.** Magnitude of PVH$^{AVP}$ inhibition during the fast inhibition phase. P=0.06 by unpaired t-test.

**g.** (Top) Mean z-score of PVH$^{AVP}$ GCaMP fluorescence during 45 minutes of recording following intraperitoneal injection of 2 M NaCl in young (3-4 mo) and aged (24-26 mo) male mice. (Bottom) Heatmaps of individual z-score traces. (Right) Area under the curve of GCaMP fluorescence z scores during 45 minutes following 2 M NaCl injection. ns., non-significant by unpaired t-test.

**h.** (Top) Mean z-score of PVH$^{AVP}$ GCaMP fluorescence during 45 minutes of recording following subcutaneous injection of AngII in young (3-5 mo) and aged (24-26 mo) male mice. (Bottom) Heatmaps of individual z-score traces. (Right) Area under the curve of GCaMP fluorescence z scores during 45 minutes following AngII injection. **P<0.01 by unpaired t-test.

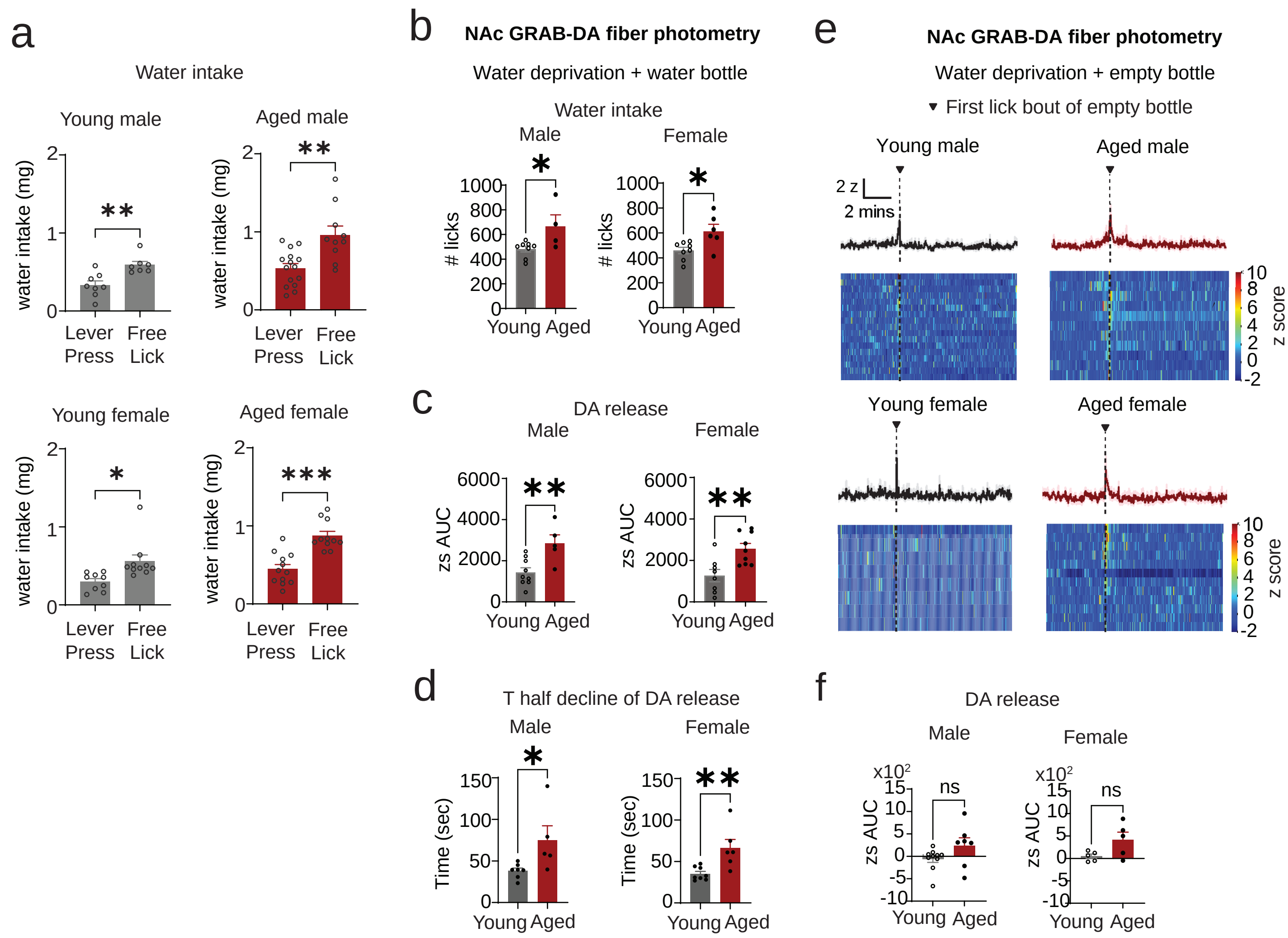


**Extended Data Fig 7. Water intake during operant and free-lick behaviors and GRAB-DA recordings during empty-bottle licking**

**a.** Water intake during 30 minutes of a fixed-ratio 1 (FR1) lever press operant task or a free lick session in males (top) and females (bottom). *P<0.05, **P<0.01 and ***P<0.001 by unpaired t-test.

**b.** Total number of water licks during 25 minutes following 24 hour water deprivation. n.s., non-significant and **P<0.01 by unpaired t-test.

**c.** Area under the curve of z-score of GRAB-DA fluorescence during the first 3 minutes following the initial water lick bout after 24 hours of water deprivation. ns, non-significant by unpaired t-test.

**d.** Time for GRAB-DA fluorescence to decay from peak to the half-max during water intake following 24 hours of water deprivation. *P<0.05 and **P<0.01 by unpaired t-test.

**e.** GRAB-DA fiber photometry recording during licking from empty bottle behavior after 24 hour water deprivation. (Top) Cumulative number of licks (blue) and mean z-score trace of GRAB-DA fluorescence in young (4-6 mo, black) and aged (24-29 mo, magenta) male and female mice, aligned to the first lick bout from empty bottle (blue dotted line). (Bottom) Heatmaps of individual z-score traces, aligned to the first lick bout from empty bottle (black dotted line).

**f.** Area under the curve of z-score of GRAB-DA fluorescence during the first 1 minute following the first lick bout from empty bottle after 24 hours water deprivation. ns, non-significant by unpaired t-test.

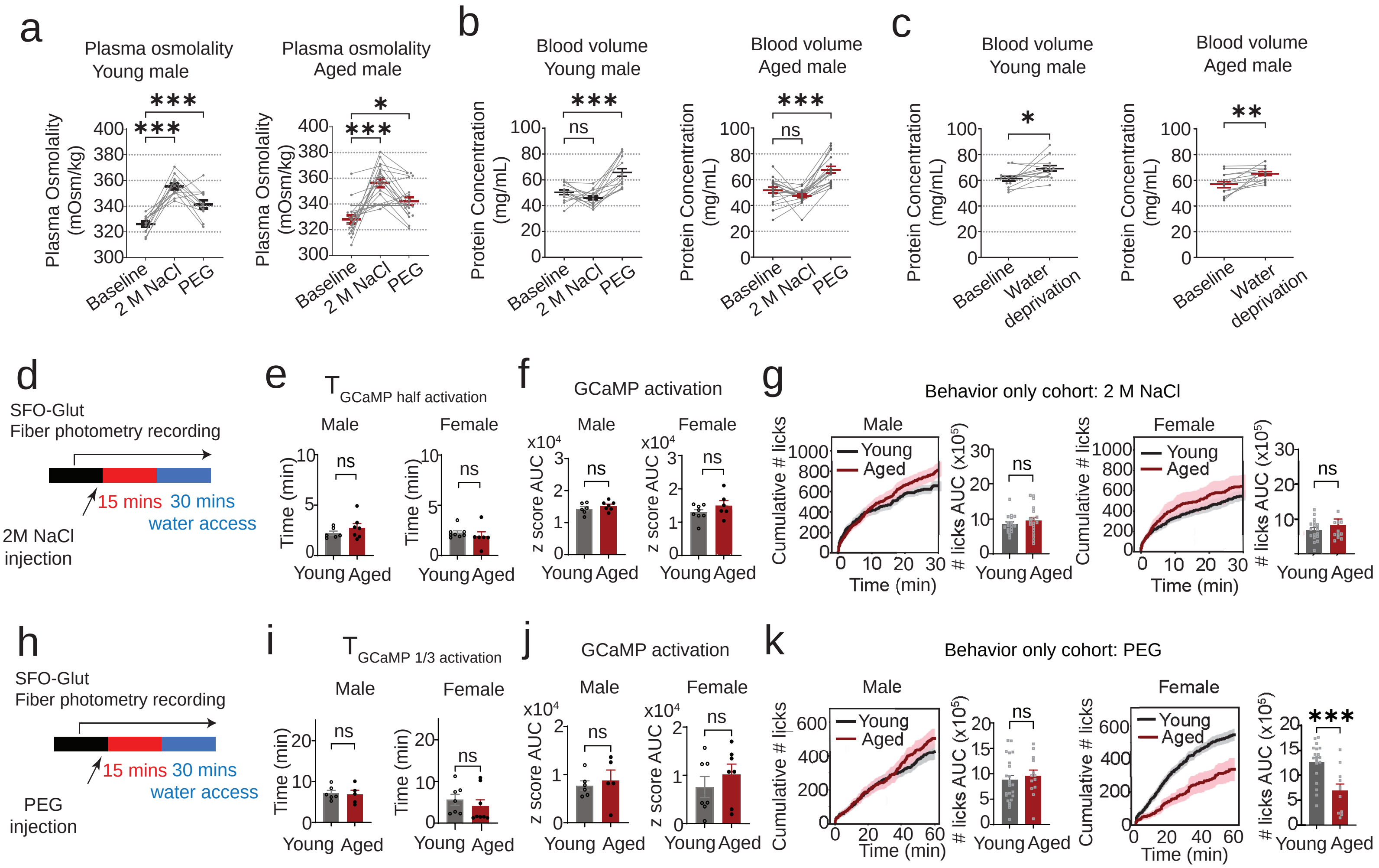


**Extended Data Fig 8. Blood parameters and SFO$^{Glut}$ activity in response to hyperosmotic and hypovolemic thirst stimuli**

**a.** Plasma osmolality in young (2-4 mo) and aged (24-25 mo) male mice in the baseline, 15 minutes after 2 M NaCl injection and 30 minutes after PEG injection. *P<0.05 and ***P<0.001 by one-way ANOVA with Dunnet's multiple comparison test.

**b.** Blood volume measured by plasma protein concentration in young (2-4 mo) and aged (24-25 mo) male mice in the baseline, 15 minutes after 2 M NaCl injection and 30 minutes after PEG injection. ns, non-significant and ***P<0.001 by one-way ANOVA with Dunnett's multiple comparison test.

**c.** Blood volume measured by plasma protein concentration in young (3-5 mo) and aged (26-27 mo) male at baseline and after 24 hours of water deprivation. *P<0.05 and **P<0.01 by paired t-test.

**d.** Experimental paradigm for SFO$^{Glut}$ fiber photometry recording during water intake after 2 M NaCl injection.

**e.** Time for GCaMP signals to rise from baseline to the half peak z-score of GCaMP fluorescence after 2 M NaCl injection. ns, non-significant by unpaired t-test.

**f.** Area under the curve of GCaMP fluorescence z scores of 30 minutes from 2 M NaCl injection to water access. ns, non-significant by unpaired t-test.

**g.** (Left) Cumulative water intake and area under the curve (AUC) of cumulative licks following 2 M NaCl injection in young (2–5 months) and aged (26–27 months) male (left) and female (right) mice ns, non-significant by unpaired t-test.

**h.** Experimental paradigm for SFO$^{Glut}$ fiber photometry recording during water intake after PEG injection.

**i.** Time for GCaMP signals to rise from baseline to the 1/3 peak z-score of GCaMP fluorescenc after PEG injection. ns, non-significant by unpaired t-test.

**j.** Area under the curve of GCaMP fluorescence z scores of 30 minutes from PEG injection to water access. ns, non-significant by unpaired t-test.

**k.** (Left) Cumulative water intake and area under the curve (AUC) of cumulative licks following PEG injection in young (2–5 months) and aged (26–27 months) male (left) and female (right) mice. ns, non-significant and ***P<0.001 by unpaired t-test.
